# Hierarchical and Spatial Mapping of Whole-Brain c-Fos Activity Reveals Distinct Opioid and Withdrawal Neuronal Ensembles

**DOI:** 10.1101/2025.09.21.677198

**Authors:** Kentaro K Ishii, Yizhe Zhang, Lynn Ren, Alex De Lecea, Srishti Bakshi, Ariyanna Haygood, Sakura Chujo, Hannah P. Stevenson, Evan Schwartz, Zhe Charles Zhou, Eric R Szelenyi, Sam A Golden, Daniel Kessler, Garret D. Stuber

**Affiliations:** Center for the Neurobiology of Addiction, Pain, and Emotion, University of Washington, Seattle, WA, 98195, USA; Department of Anesthesiology and Pain Medicine, University of Washington, Seattle, WA, 98195, USA; Department of Pharmacology, University of Washington, Seattle, WA, 98195, USA; Department of Neurobiology and Biophysics, University of Washington, Seattle, WA, 98195, USA; Department of Statistics and Operations Research, University of North Carolina at Chapel Hill, Chapel Hill, NC, 27599, USA; School of Data Science and Society, University of North Carolina at Chapel Hill, Chapel Hill, NC, 27599, USA

## Abstract

How opioid exposure and withdrawal states shape brain activity at the systems and circuit level remains poorly understood. Here, we use whole-brain, cellular-resolution c-Fos mapping to define brain-wide activity patterns and neuronal ensembles associated with morphine administration and withdrawal. To account for the brain’s anatomically nested structure, we developed and applied a hierarchical statistical framework that detects region-specific changes in activity and outperforms conventional methods that treat brain regions as individual, unrelated units. These distributed signals formed ensembles with consistent and anatomically structured patterns of activity, both within subregions and across multiple connected brain areas. By combining TRAP2-based activity tagging with acute whole-brain c-Fos staining, we identified morphine- and withdrawal-activated ensembles and found that they are largely non-overlapping at the single-cell level, even within the same brain region. Integration with existing spatial transcriptomics datasets identified molecular markers for these state-specific ensembles in key brain areas such as the nucleus of accumbens, amygdala and ventral tegmental areas. Lastly, by integrating Allen mouse whole-brain transcriptional datasets, we identified the molecular identity of the morphine- administration and withdrawal ensembles. These findings define dissociable neuronal ensembles that encode opposing drug states and introduce a scalable framework for linking whole-brain activity to molecular and circuit-level mechanisms.

**Highlights:**

- Whole-brain neural activity mapping identifies the regional and spatial difference of morphine-administration and withdrawal neuronal ensembles.
- BRANCH, a hierarchical statistical testing framework, offers increased sensitivity and anatomical interpretability for whole-brain datasets.
- TRAP2-based activity tagging reveals brain-wide separation of acute morphine and withdrawal ensembles at the cellular level.
- A systematic analysis of the whole brain neural activity data combined with spatial transcriptomic data revealed molecular features of opioid-related neural ensembles.

## Introduction

Opioids engage widespread neural systems that regulate reward, pain, motivation, and stress^1–3^. The transition from acute drug exposure to chronic use and withdrawal involves major shifts in brain function, including changes in the circuits that encode positive and negative motivational states^4^. These neurobiological transitions, from the rewarding and analgesic effects of initial opioid administration to tolerance with repeated use and then to the dysphoria and stress of withdrawal, are thought to drive compulsive drug seeking and high relapse risk^5,6^. Among the vast variety of opiates, morphine is most widely used in clinical settings due to its potent analgesic properties and efficacy in managing moderate to severe pain through well-characterized pharmacokinetics. At the molecular level, morphine acts primarily through μ-opioid receptors (MOR), with significantly lower affinity for κ-opioid (KOR) and δ-opioid (DOR) receptors. These are broadly distributed across the brain but exhibit region- and cell type-specific expression patterns^7^, which are considered to contribute to the complex and heterogeneous effect of opiate drugs. Given this complexity, it is essential to characterize how opioid-associated molecular signals manifest within specific neural circuits.

Historically, insights into opioid-induced neural activity have largely come from studies targeting specific brain regions within reward-related circuits. Foundational work from Berridge and colleagues has shown that the behavioral effects of opioid signaling can be organized at a fine anatomical scale, conceptualized as the opioid hedonic "hotspots–coldspots"^8–11^; a model which suggests that brain structures in the reward system contains anatomically distinct functional sub regions that encode the hedonic value of opioids. Microinjections of opioid agonists into restricted subregions of structures such as the nucleus accumbens^12^ and ventral pallidum^13^ can strongly enhance hedonic “liking” reactions, whereas injections into adjacent zones can suppress these behaviors^14^. These hedonic hotspots and coldspots reveal that opposing behavioral effects can arise from closely neighboring neuronal populations, emphasizing the need for high-resolution, spatially precise mapping to understand how opioids influence brain function. Despite considerable progress in defining the neural mechanisms of opioid action, most studies have focused on a limited number of brain regions, particularly canonical reward and aversion centers such as the nucleus accumbens, ventral tegmental area, amygdala, and periaqueductal gray. These efforts have relied heavily on hypothesis-driven approaches that cannot detect distributed or noncanonical networks. Analyses are often performed at the level of gross anatomical regions, which ignores the brain’s nested anatomical structure and can obscure fine-scale spatial patterns. Moreover, the molecular identity of neurons within these state-specific neural ensembles remains largely unknown, posing a challenge for investigating their functional properties using modern genetic tools.

Recent advances in whole-brain tissue clearing and light-sheet imaging have enabled high-throughput, cellular-resolution mapping of immediate early gene (IEG) activity^15,16^. These techniques offer a powerful means to analyze the spatial, cellular, and molecular properties of neural ensembles, owing to the highly standardized datasets they produce. Indeed, whole brain mapping of opioid-related neural activity has been conducted in several studies^17–22^, but they lacked comprehensive analysis of the spatial and cellular distribution of opioid administration and withdrawal ensembles. In this study, we introduce and apply a pipeline for hierarchical and spatial analysis of brain-wide expression of c-Fos, a widely used IEG, across multiple morphine-related states in mice. We applied a brain-wide regional analysis with control for hierarchy that leverages the brain’s anatomically nested structure to detect region-specific changes in activity with greater sensitivity and anatomical interpretability than conventional region-by-region testing. This method revealed key brain regions and hierarchical structures that are differentially engaged across morphine-related states. Further building on these findings, we mapped MOR+ cells across the brain to determine how receptor expression correlates with morphine-induced neural ensembles. We next classified neural ensembles to identify functional clusters of brain regions that co-activate during administration, withdrawal, or both. We then resolved these distributed activity patterns into anatomically structured ensembles using semi-negative normalized matrix factorization approaches and determined their cellular specificity using genetic tagging of neural ensembles combined with whole-brain c-Fos immunolabeling. Even in regions with elevated activity in both administration and withdrawal, we found that the underlying neuronal populations were largely distinct. Importantly morphine-administration and withdrawal aligned with the hotspot–coldspot framework, particularly in the nucleus accumbens. Finally, we integrated the results with existing spatial transcriptomics datasets to identify candidate molecular markers in key morphine-state encoding brain regions. This multiscale approach defines dissociable neuronal ensembles that encode opposing drug states and establishes a generalizable framework for linking brain-wide activity to molecular and circuit-level mechanisms.

## Results

### Whole brain neural activity during distinct morphine-related states

To investigate how the brain responds to distinct phases of morphine exposure and withdrawal, we performed cellular-resolution, whole-brain activity mapping using c-Fos immunolabeling combined with tissue preservation and clearing. c-Fos protein is a well-established molecular marker of increased neuronal excitability, which we use here as an approximation of neural activity. We collected brain tissue from six experimental groups (**Figure 1A**). To assess the acute response to morphine, one group received a single injection of 20 mg/kg morphine and was perfused two hours later (acute group). A control group received saline and was perfused on the same schedule (saline group). To examine the effects of repeated drug exposure, a chronic group received twice-daily injections for seven days with escalating doses from 0.1 to 20 mg/kg, followed on the eighth day by 20 mg/kg morphine and perfusion two hours later. To model early spontaneous withdrawal, another group received the same escalating morphine schedule but saline on the final day (earlyWD group). To examine the impact of drug re-exposure after abstinence, a re-exposure group underwent the same seven-day morphine regimen, remained drug-free for 21 days, then received 20 mg/kg morphine and was perfused two hours later. A late withdrawal group followed the same schedule but received saline on the final day (lateWD group).

**Figure 1:**
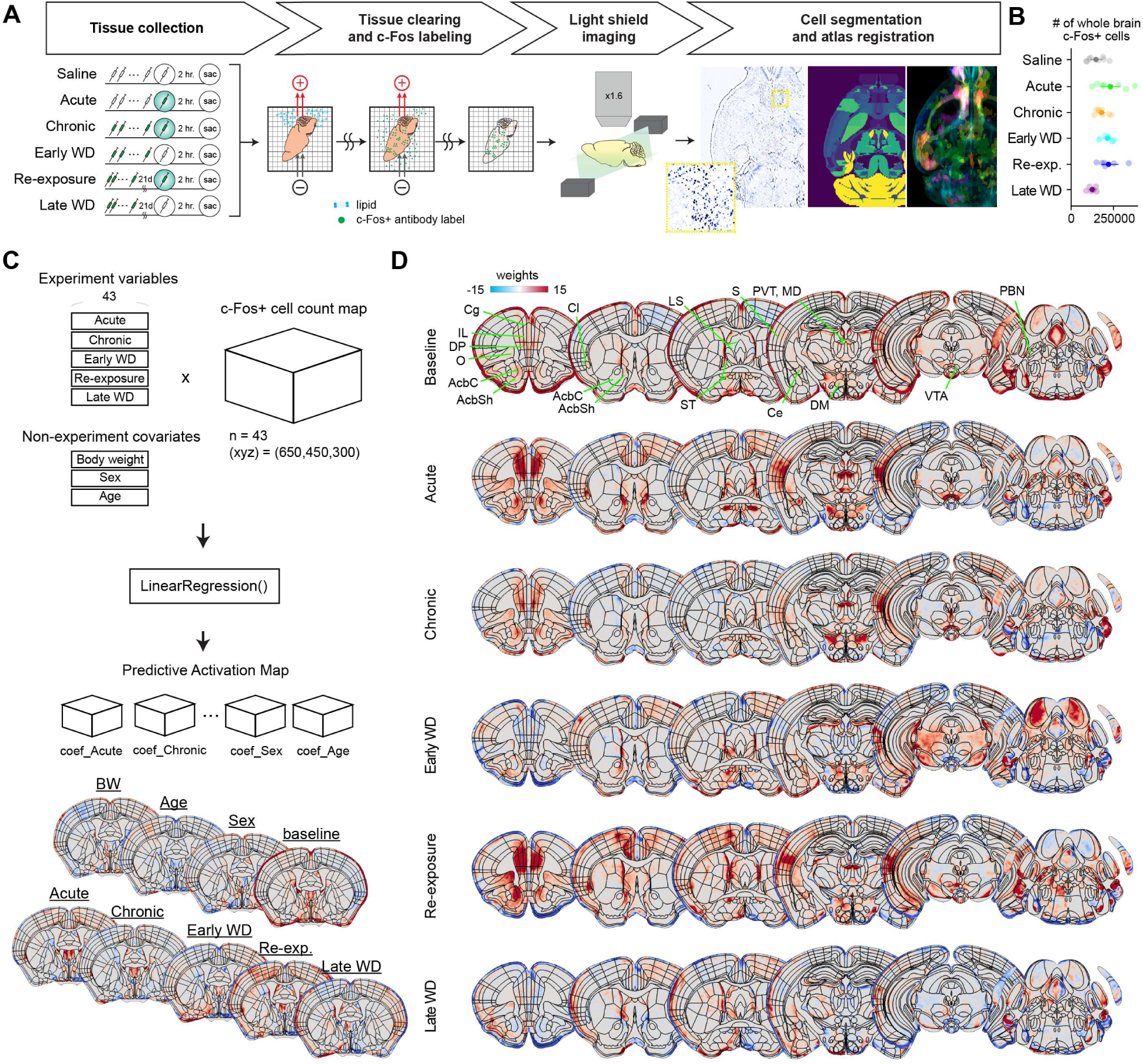
Whole-brain analysis of neural response to morphine administration and withdrawal. (A) Overview of the experiment pipeline. To model different time points during morphine treatment, we prepared 5 conditions with different drug schedules. Two hours after the final drug administration, animals were perfused to collect brain tissue. Brains were first processed in SHIELD for preservation, then cleared and stained for c-Fos. Tissues were then imaged on a light sheet microscope. (B) Total number of c-Fos+ cells in each condition. (C) Regression analysis to correct for non-experiment variables. Experiment variables (5 drug conditions, saline as baseline) as well as non-experimental covariates (body weight, age, sex) were used to regress c-Fos+ cell count map (x,y,z) = (650,450,300) from n = 44 animals. The beta coefficient from the regression were used as condition-specific predictive activation score relative to saline. (D) Representative 50-μm coronal planes showing predictive activation map for each drug condition. Saline: n = 8, Acute: n = 7, Chronic: n = 8, EarlyWD: n = 8, Re-exposure: n = 6, LateWD: n = 6.

We processed brains with tissue preservation reagents^23^, followed by active delipidation and antibody staining. Samples were imaged using light-sheet microscopy, registered to the Unified Mouse Brain Atlas^24^, and analyzed with a high-performance computing pipeline to quantify c-Fos–positive cell counts by region and in voxel space. On average, brains contained 1.8 ± 0.6 × 10⁵ c-Fos–positive cells (mean ± SEM), with saline and late withdrawal groups showing fewer counts than other conditions (**Figure 1B**). We then performed regression analysis on voxelated c-Fos data to generate predictive activation maps, which reflect changes in c-Fos counts relative to baseline while controlling for non-experimental variables (**Figures 1C and 1D**). This analysis revealed widespread activation, most prominently in ventral striatum, prefrontal cortical regions, and the amygdala. To determine which of these activity changes were statistically robust and anatomically interpretable across the brain’s nested structure, we next applied a statistical testing procedure tailored to whole-brain datasets to compare morphine and withdrawal states.

### BRANCH testing identifies hierarchical brain regions engaged across morphine-related states

To systematically identify brain regions engaged across distinct stages of morphine treatment, we quantified c-Fos–positive (c-Fos+) cell density across the entire brain and developed <u>b</u>rain-wide regional <u>an</u>alysis with <u>c</u>ontrol for <u>h</u>ierarchy (BRANCH), a hierarchical statistical testing procedure based on the TreeBH error-controlling method^25^ and extends it to leverage the anatomically nested organization of the brain atlas (**Figure 2A**). Traditional whole-brain analyses typically treat regions as unrelated units, applying multiple comparison correction to a flat list of regions without regard to structural granularity. This approach discards important anatomical information, obscures dependencies among related subregions, and can fail to detect divergent effects within structural families. For example, when opposing signals arise in different subregions of a region (e.g., one subregion increases and another decreases), flat-list testing may erroneously conclude there is no effect, whereas a tree-aware method can identify such cancellation patterns as meaningful.

**Figure 2.**
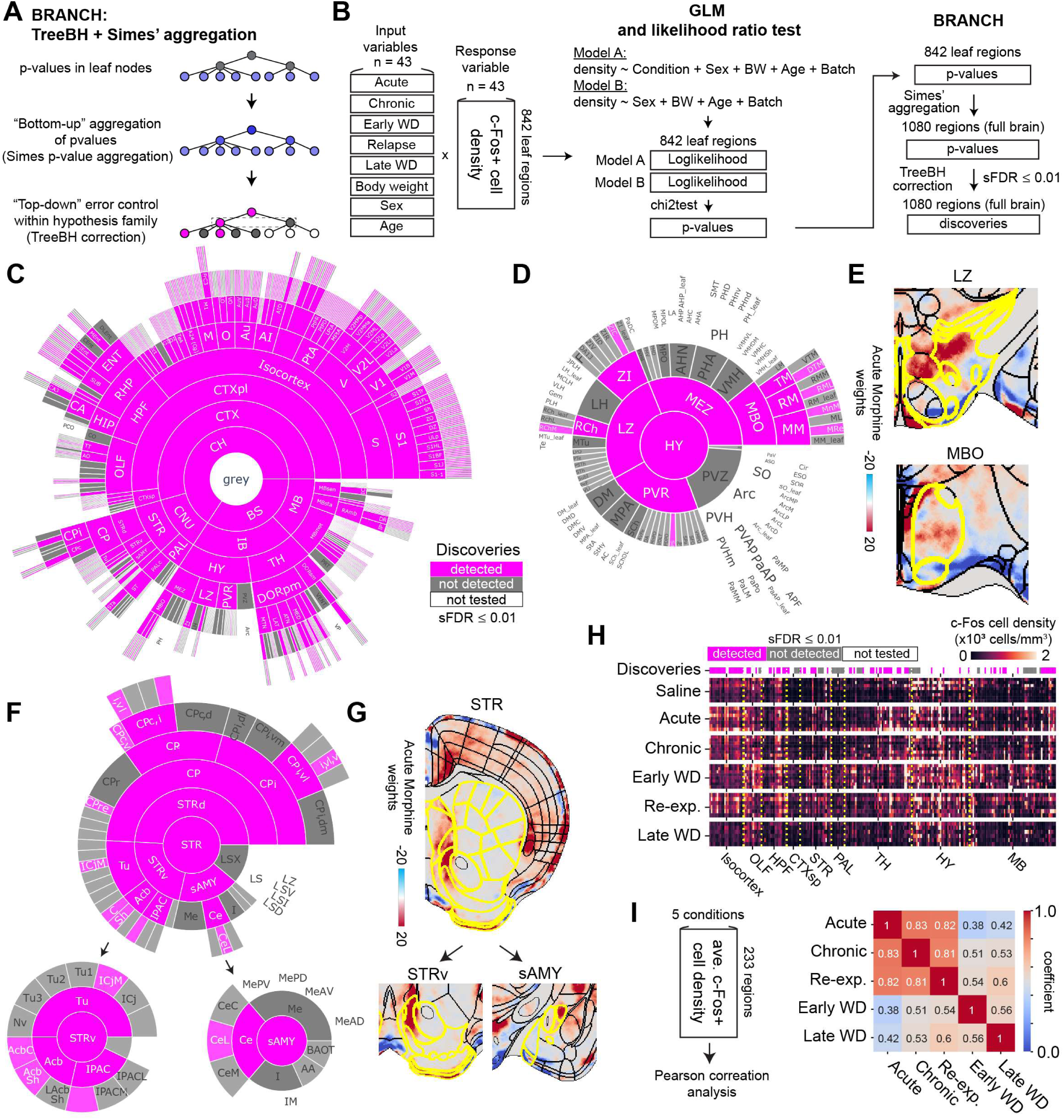
Usage of BRANCH procedure for robust hierarchical statistical testing in whole-brain data. (A) Schematic showing the steps of BRANCH error control procedure. (B) Schematic illustration of statistical analysis. To evaluate the effect of morphine administration, a GLM and likelihood ratio test was conducted on 842 leaf nodes of the Unified mouse brain atlas ontology tree. P-values generated in the GLM and likelihood ratio test were further processed using BRANCH. (C) Sunburst plot showing statistical testing results across the ontology. Purple: rejected null hypothesis; gray: not rejected; white: not tested. (D) Detailed analysis of the hypothalamus subtree. (E) Representative 50-μm coronal planes showing predictive activation map of acute morphine condition in the lateral zone of hypothalamus (LZ) and mammillary body (MBO). (F) Detailed analysis of the striatal subtree. (G) Representative 50-μm coronal planes showing predictive activation maps of acute morphine condition in the ventral striatum (STRv) and striatal amygdalar area (sAMY). (H) Heatmap showing c-Fos+ cell density for each region in a 233 pre-selected list of brain regions (**Table 1**). Discoveries for each brain region are shown above. (I) Correlation matrix using the average c-Fos+ cell density data for 233 pre-selected brain regions for each drug conditions. Saline: n = 8, Acute: n = 7, Chronic: n = 8, EarlyWD: n = 8, Re-exposure: n = 6, LateWD: n = 6. Additional statistics are shown in **Table 2**.

At its core, BRANCH consists of two phases: 1) “bottom-up” aggregation of p-values and 2) “top-down” error control in a tree-structured set of hypotheses defined by an annotated brain atlas. Each brain region is represented as a node, with subregions as children and higher-order structures as parents. We first assigned p-values to the 842 terminal (leaf) regions of the atlas. In the first part of BRANCH, the p-values for all nodes are obtained by recursively aggregating among its children using Simes’ method, which essentially tests for deviations from the null among descendants. In the second part, selective FDR (sFDR) control was applied recursively using TreeBH^25^, a top-down pruning algorithm. Nodes were tested only if their parent node was rejected and the local q-value threshold (sFDR < 0.01) was met.

To assess morphine effects on neural activity, we used a generalized linear modeling (GLM) framework to compare c-Fos+ cell density across experimental conditions while adjusting for biological covariates (**Figure 2B**). For each brain region, we fit a full model including all morphine treatment conditions (Acute, Chronic, Re-exposure, Early Withdrawal, Late Withdrawal) and covariates such as sex, body weight, and age. We then fit a reduced model excluding treatment conditions and used a likelihood ratio test (LRT) to determine whether inclusion of treatment improved model fit. The resulting p-values were input to the BRANCH, which identified significant discoveries distributed throughout the brain (**Figure 2C**).

First, we found a broad distribution of significant discoveries in the cortical regions, particularly the isocortex (199 of 303 significant leaf regions). In contrast, we detected sparse signals in the hypothalamus, with significant effects in only 7 of 98 leaf regions: retro mammillary nucleus, lateral part (RML); medial mammillary nucleus, median part (MnM); and mammillary recess of the third ventricle (MRe) (**Figures 2D and 2E**). In the striatum (STR) subtree, we identified 10 of 63 significant regions (**Figures 2F and 2G**). Within the ventral striatum (STRv), significant effects were present in all subregions, including the accumbens (Acb), olfactory tubercle (Tu), and interstitial nucleus of the posterior limb of the anterior commissure (IPAC). In the Acb, a localized neural activity signal was detected in both the core (AcbC) and shell (AcbSh) but not in the lateral shell (LAcbSh). In striatal amygdala (sAMY) regions, significant effects were confined to the central amygdala (Ce), typically in the lateral part (CeL). These findings demonstrate how BRANCH controls multiple comparison error while preserving the hierarchical structure of the atlas.

For further analysis of by-region neural activity data, we decided to focus on a predefined set of 233 disjoint brain regions (i.e., no parent–child relationships). These regions were chosen unbiasedly throughout the whole-brain and matched the resolution of standard neuroscience research (**Table 1**). Here, we first found that 89 of these 233 regions had a significant change in neural activity in morphine-related conditions (**Figure 2H**). To evaluate the impact of BRANCH, we compared it to two flat multiple testing approaches (**Figure S1A**). In the most classical method (which we denote FlatBH) method, p-values were computed from average c-Fos+ density per region and corrected using Benjamini–Hochberg correction. In the FlatBH with Simes’ variant, leaf-region p-values were aggregated with Simes’ method before Benjamini–Hochberg correction. Note that BRANCH combines Simes’-based aggregation with TreeBH hierarchical correction. Both FlatBH with Simes’ aggregation and BRANCH improved statistical power compared to the non-Simes approach (**Figure S1)**. Regions identified using Simes-based p-value aggregation had significantly more child nodes, suggesting that ignoring subregion data may miss a substantial fraction of true effects. Simulations using randomly selected node sets confirmed these findings were not specific to our region selection (**Figures S1I–J**).

To examine relationships among morphine exposure conditions, we calculated pairwise Pearson correlations of average c-Fos+ density across regions. Acute, Chronic, and Re-exposure conditions were highly correlated (r > 0.8), consistent with shared neural activation patterns (**Figure 2I).** Withdrawal conditions (EarlyWD, LateWD) were less correlated with morphine administration groups and with each other. Together, these results identify a brain-wide distributed set of regions containing morphine administration–related and withdrawal-related signals and demonstrate that BRANCH improves both sensitivity and anatomical interpretability in whole-brain activity mapping.

### Increased neural activity after acute morphine administration

Acute morphine exposure produced widespread increases in neural activity across the brain (**Figure 3A and Figure S2**). To fully evaluate the specific brain regions that are immediately affected by morphine, we compared the change in c-Fos cell density between saline and acute morphine conditions. Among the 89 brain regions identified by BRANCH as having differential activity across conditions, 31 regions showed a significant increase in c-Fos density after acute morphine administration relative to saline (**Figure 3A and 3B; Figure S5B**). Surprisingly, the claustrum (Cl) was the region with the largest difference in c-Fos density (c-Fos cell counts normalized by region size). From the voxel-based analysis, we observed that the changes within Cl were concentrated in both dorsal and ventral subregions and throughout the AP axis (**Figure S2**). Similarly, we observed that most prefrontal cortical regions (prelimbic (PrL), infralimbic (IL), cingulate (Cg), dorsal peduncular (DP), orbitofrontal (O)) were strongly activated by acute morphine administration. We also observed signals in the AcbC and AcbSh, as suggested by the region-based analysis, and consistent with previous literature^8,26,27^. Within the thalamic regions, we found concentrated signals in most of the medial thalamic regions such as mediodorsal (MD), parataenial (PT), paraventricular nucleus (PVT), central medial (CM), intermediodorsal (IMD), posteromedian (PoMn) and interanterodorsal (IAD) thalamic region. In the subcortical regions, we found increased neural activity in the amygdala and striatal amygdala structures, such as the central (Ce), lateral (La) and basolateral (BLA) amygdala. In the caudate putamen structure, we observed signals in the rostral-extreme part (CPre) and in the caudal part (CPc), especially in its caudal extreme subregion (CPce) structure (also known as the tail of the striatum). In contrast, the hypothalamus showed minimal activation. No hypothalamic leaf nodes were significantly enriched for c-Fos+ cells, although voxel maps revealed localized activity within the lateral hypothalamic area (LH) near the dorsomedial hypothalamus (DM). This suggests that morphine’s acute activation of hypothalamic circuits is spatially restricted and may not align with traditional atlas boundaries. Other regions where we observed increased c-Fos expression were the tegmental areas such as the ventral tegmental area (VTA) and subpeduncular tegmental nucleus (SPTg). In midbrain, we found signals concentrated in neuromodulator hubs such as in the dorso raphe (DR), the ventrolateral part of the parabrachial nucleus (PB), and the locus coeruleus (LC). In sum, acute morphine administration recruits a distributed set of cortical, striatal, thalamic, and brainstem regions with distinct spatial patterns. It engages medial thalamic and amygdalar hubs and largely spares classical hypothalamic circuits. These patterns highlight a rapid, systems-level reorganization of forebrain and midbrain networks.

**Figure 3.**
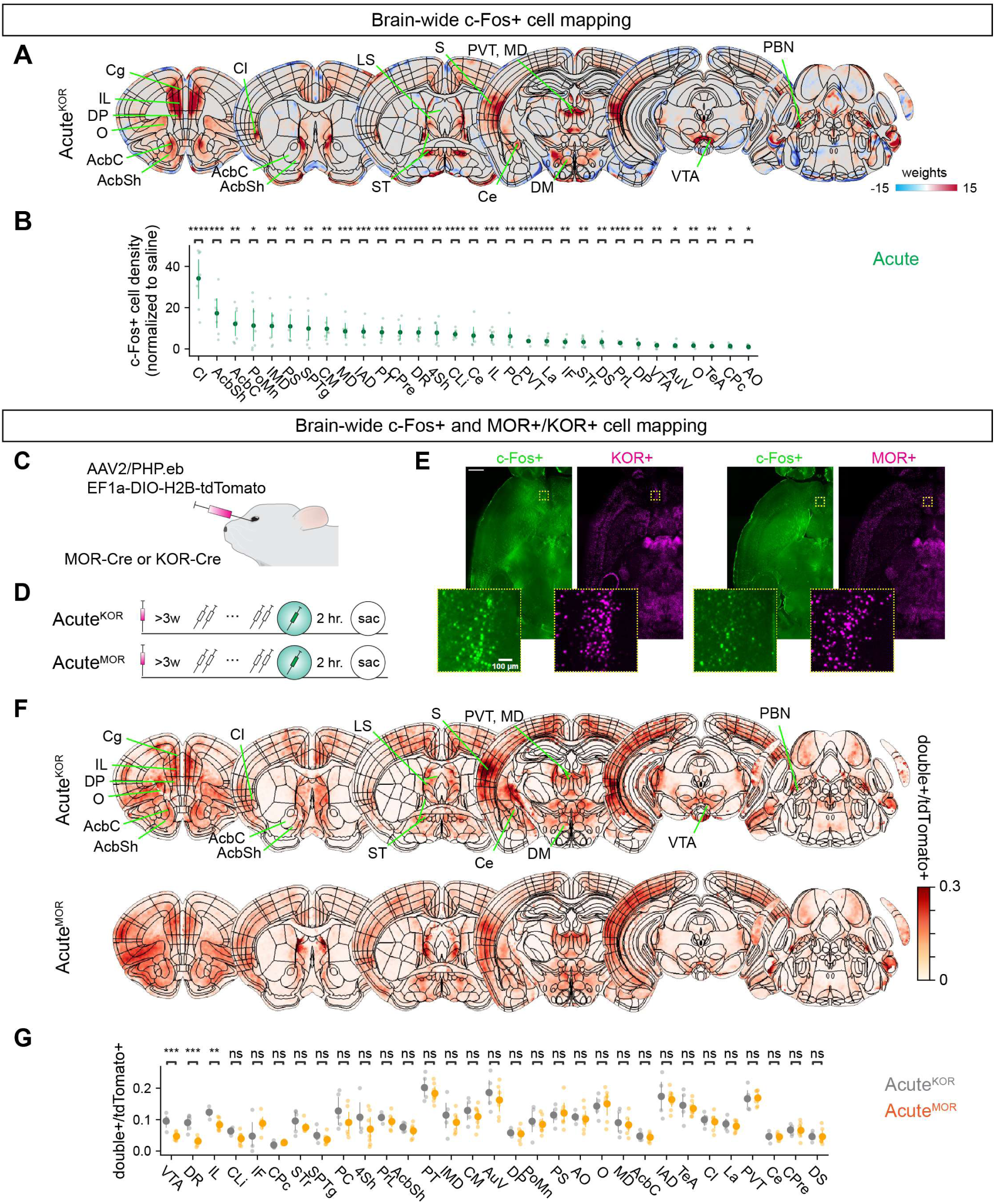
Increased neural activity after acute morphine administration. (A) Predictive activity map for acute morphine condition. Detailed visualization is shown in **Figure S2**. (B) Scatter plot showing brain regions with significant difference in c-Fos+ cell density in *post hoc* testing of acute morphine vs saline conditions among the 89 brain regions identified using BRANCH. Student’s t-test with Benjamini/Hochberg correction (alpha = 0.05). Full results are shown in **Figure S5B**. Additional statistics are shown in **Table 3**. Saline: n = 8, Acute: n = 7. (C) Schematic illustration showing retroorbital injections to label KOR/MOR+ cells. AAV2.PHP.eb EF1a-DIO-H2B-tdTomato was injected retro-orbitally into KOR-Cre+ or MOR-Cre animals. (D) Overview of experiment design. More than two weeks after retroorbital injection, animals went through acute morphine drug schedule. (E) c-Fos+ and tdTomato+ cells were registered to the Unified mouse brain atlas for further quantification. (F) Representative 50-μm coronal planes showing average double+ cells normalized by the total number of tdTomato+ cells. (G) Scatter plot showing double+ cells normalized by the total number of tdTomato+ cells among the 31 brain regions that were identified as acute morphine responsive regions in Figure 3B. Student’s t-test with post-hoc Benjamini/Hochberg correction over brain regions. Acute^KOR^: n = 5. Acute^MOR^: n = 7. Additional statistics are shown in **Table 4**.

### Mu-opioid receptor cells are selectively suppressed by morphine in a region-specific manner

The preceding analyses characterized brain-wide patterns of activity in response to acute morphine, but they do not directly address how receptor expression constrains the recruitment of these ensembles. Opioid effects are mediated primarily by Gi/o-coupled opioid receptors, with morphine showing high efficacy at the μ-opioid receptor (MOR) and much lower efficacy at the κ-opioid receptor (KOR). Both MOR and KOR activation suppresses neuronal excitability by inhibiting transmitter release and promoting hyperpolarization, and morphine-induced increases in neural activity are often attributed to circuit-level disinhibition. In the ventral tegmental area (VTA), for example, MORs are enriched in inhibitory neurons both within and outside the VTA^20,28–31^, hypothesized to produce disinhibition of VTA dopamine (DA) neurons. Whether this mechanism generalizes to other brain regions remains unresolved. To test how receptor distribution relates to morphine-induced activity at the whole-brain level, we selectively labeled MOR- and KOR-expressing neurons by retro-orbital injection of AAV-PHP.eB driving Cre-dependent nuclear tdTomato in MOR-Cre or KOR-Cre mice (**Figure 3C**). After allowing >3 weeks for expression, animals were administered acute morphine and perfused 2 hours later for brain clearing and whole-brain c-Fos immunolabeling (acute^MOR^ and acute^KOR^ groups) (**Figure 3D**). Cleared brains were imaged via light-sheet microscopy to identify c-Fos+, tdTomato+, and double-labeled (double+) neurons (**Figure 3E**), allowing a direct comparison of morphine-induced activation in a MOR⁺ population versus a less-sensitive KOR⁺ population.

Both KOR+ cells and MOR+ cells were widely distributed across the brain, with MOR+ cells being more dominant using our methods (**Figure S3A** and **S3B**) (whole-brain KOR+ cells, 2.1 ± 0.7 × 10^6^ cells, n = 5. MOR+ cells, 3.8 ± 1.0 × 10^6^ cells, n = 7. Mean ± SEM). Spatial mapping further revealed that double+ cells normalized by the total tdTomato+ cells (double+/tdTomato+) in acute^MOR^ and acute^KOR^ groups resemble the acute morphine predictive activity map (**Figure 3E**). Among the 31 brain regions that had significant c-Fos activity in acute morphine conditions (**Figure 3B**), we found that only 3 regions showed significant differences in double+/tdTomato+ between acute^KOR^ and acute^MOR^: DR, VTA, and IL (**Figure 3F**). Region-specific anterior-posterior mapping showed that KOR+ c-Fos+ cells were enriched in the vlPAG and DR (**Figure S3C–F**). In contrast, MOR+ c-Fos+ cells had weaker signals in these region, despite having similar amount of tdTomato+ cells. KOR+ c-Fos+ cells but not MOR+ c-Fos+ cells were enriched in the VTA, particularly in the parabranchial pigmented nucleus subregion **(Figure S3G–S3J**). These findings indicate that morphine’s inhibitory effect toward MOR+ cells occur in a region-specific manner. Furthermore, the widespread morphine-induced activation seen in regions such as the ventral striatum, claustrum, extended amygdala, and medial thalamus is unlikely a result of direct MOR action within those regions, instead reflects indirect recruitment of a broader circuit mechanisms and/or local modulation, such as at the presynaptic terminal, which does not induce somatic changes.

### Morphine administration and withdrawal drive divergent brain-wide activity patterns

Chronic morphine exposure produces tolerance and dependence, and withdrawal elicits stress-related and aversive states^32^. To identify the brain-wide activity patterns underlying these contrasting states, we directly compared chronic morphine (chronic) and early spontaneous withdrawal (earlyWD) conditions (**Figure 4A**). Spatial mapping of modeled c-Fos activity revealed clear regional differences between the two conditions (**Figure 4B and Figure S4A**). Prefrontal cortical regions such as IL and DP, and medial thalamic regions such as PVT and MD showed larger increases in c-Fos cells in chronic morphine conditions, while the Ce showed increases in the earlyWD conditions. These regions are suggested to be involved in pain, threat, and stress processing^33–35^; this aligns with our finding of their recruitment during withdrawal^35,36^. To systematically classify brain regions based on their response profiles across chronic morphine and earlyWD conditions, we applied a dimensionality reduction and clustering framework (**Figure 4C–F**). We used Uniform Manifold Approximation and Projection^37^ (UMAP) to embed the z-scored c-Fos density changes from chronic and earlyWD conditions for 89 brain regions identified by BRANCH in **Figure 2**. In this embedding, Euclidean distances reflect similarity in activity patterns between regions (**Figure 4C and 4D; Figure S5B**). Agglomerative hierarchical clustering of the UMAP coordinates yielded four distinct functional clusters. Interestingly, these clusters mapped closely to major anatomical systems. Cluster 1 grouped medial thalamic regions together with granular (GI), dysgranular (DI), and agranular (AI) subregions of the insular cortex, Acb and the claustrum (Cl), suggesting a possible network for integrating interoceptive and motivational state information. Cluster 2 contained predominantly prefrontal cortical regions, while cluster 3 enriched limbic–amygdalar structures, including Ce and La subdivisions. Three clusters 1, 2 and 4, had larger z-scored c-Fos density in chronic conditions than in earlyWD conditions, indicating that these networks, although anatomically distinct, share a bias toward chronic morphine engagement (**Figure 4E**). In contrast, cluster 3 had larger average scaled density in earlyWD conditions, consistent with a coordinated role for limbic–amygdalar regions in withdrawal-related processes.

**Figure 4.**
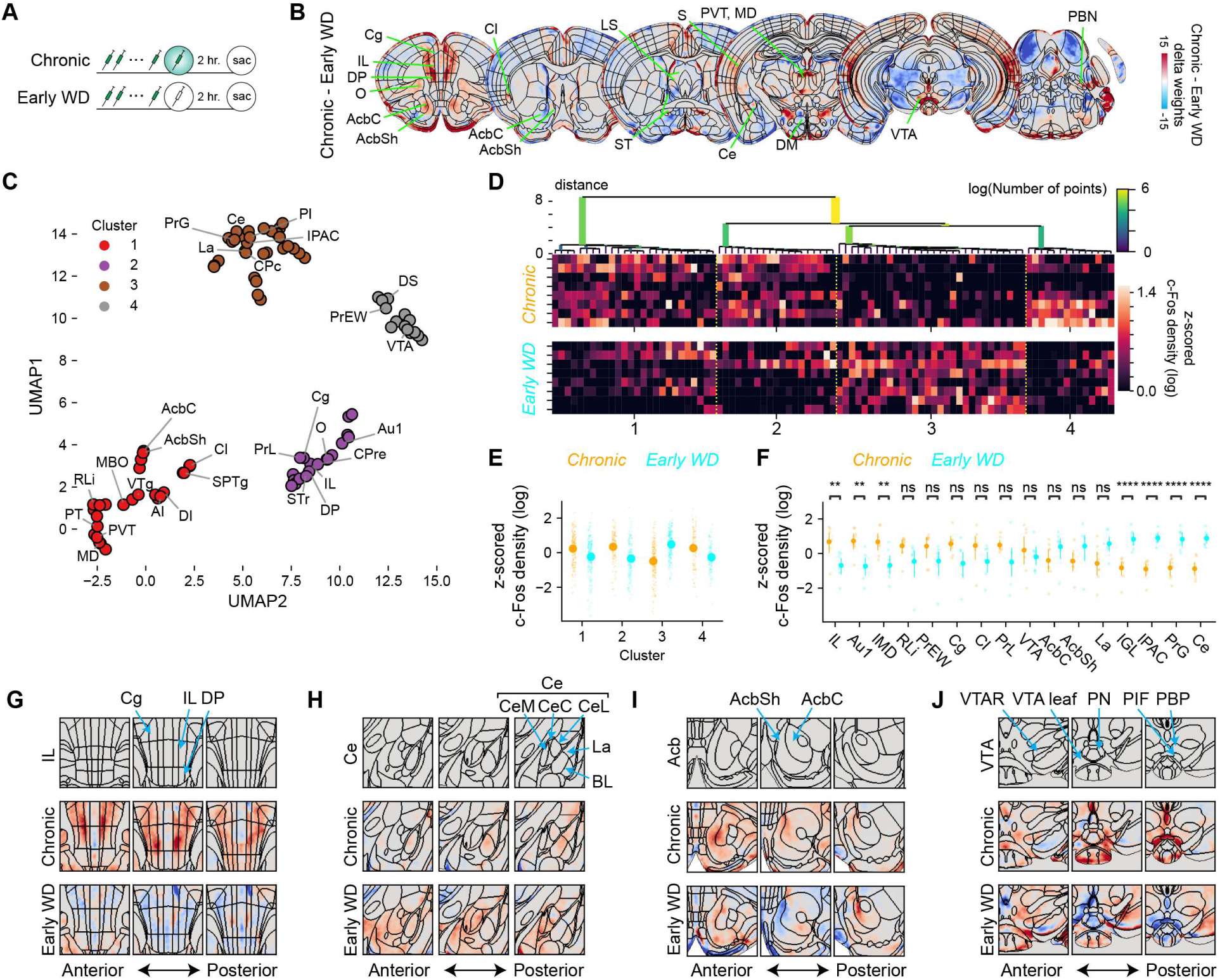
Distinct neural activity pattern during chronic morphine administration and early withdrawal. (A) Schematic showing drug schedule for chronic morphine and early withdrawal conditions. (B) Representative 50-μm coronal planes showing difference in predictive activity map between chronic and earlyWD. Detailed visualization is shown in **Figure S4A**. (C) Uniform Manifold Approximation and Projection (UMAP) display of c-Fos+ cell density data from 95 brain regions. Colors indicate clusters identified by HDBSCAN. (D) Heatmap showing the z-scored c-Fos density across regions within each identified cluster. Each row corresponds to a subject, top: Chronic morphine; bottom: Early Withdrawal. Hierarchical lineage tree generated by HDBSCAN is shown on top. (E) Scatter plot showing z-scored c-Fos density for brain regions in each cluster (Cluster 1: n = 208, EarlyWD n = 208. Cluster 2: Chronic n = 152, EarlyWD n = 152. Cluster 3: Chronic n = 240, EarlyWD n = 240. Cluster 4: Chronic n = 112, EarlyWD n = 112.). (F) Scatter plot showing *post hoc* analysis of group differences in z-scored c-Fos density in selected brain regions from among those identified by BRANCH (full results are shown in **Figure S4B**). Student’s t-test with Benjamini/Hochberg correction (alpha = 0.05). Chronic: n = 8, EarlyWD: n = 8. Additional statistics are shown in **Table 4**. (G–J) Representative 50-μm coronal planes showing predictive activity map in the prefrontal regions (G), amygdalar regions (H), nucleus of accumbens (I) and ventral tegmental area (J). VTA leaf refers to subregions of VTA that were not categorized as other subregions (VTAR: ventral tegmental area rostral. PN: paranigral nucleus. PIF: parainfudibular nucleus. PBP: parabrachial pigmented nucleus.).

At the level of individual regions, IL, primary auditory cortex (Au1) and IMD had significantly greater activity in chronic morphine, whereas Ce, pregeniculate nucleus (PrG), interstitial nucleus of the posterior limb of the anterior commissure (IPAC) and intrageniculate leaflet (IGL) were more active in earlyWD (**Figure 4F and Figure S4B**). When we further examined the spatial pattern of modeled activity maps, we observed increased neural activity in IL and amygdalar regions throughout the anterior-posterior axis in the chronic morphine and earlyWD conditions, respectively (**Figure 4G and 4H**). In contrast, key brain regions in the reward system such as the VTA, AcbC and AcbSh showed no significant difference at the region level. However, we observed unique patterns of activation in the spatial distribution. In the Acb, chronic morphine administration increased neural activity in the anterior part, while earlyWD increased neural activity in the posterior part (**Figure 4I**). In the VTA, chronic morphine administration increased neural activity throughout the anterior-posterior axis, but in the posterior part within the parainfudibular nucleus (PIF), while earlyWD increased neural activity in the VTA rostral subregion (VTAR), but not in the posterior parts (**Figure 4J**). These results suggested that even in brain regions with no significant difference at the region level, they can show segregated increases in neural activity between conditions.

Together, our region-level analysis shows that chronic morphine administration and early withdrawal recruit largely distinct sets of brain regions, with key differences in prefrontal–thalamic versus limbic–amygdalar engagement. However, the results also indicated that further spatial evaluation was necessary.

### Morphine administration and withdrawal recruit spatially distinct neural activity patterns

To systematically uncover structured patterns of brain-wide activation across experimental groups, we applied a semi-nonnegative matrix factorization (semi-NMF) pipeline^38^ to c-Fos+ cell count maps (**Figure 5A**). Briefly, c-Fos+ cell count maps were downsampled and decomposed into 22 latent spatial factors and corresponding subject-specific loading weights using a sparse Poisson semi-NMF model (**Figure 5B**, **Figure S6A**). Each factor represented a spatially resolved ensemble, and the corresponding weights reflected its contribution to each animal’s whole-brain c-Fos+ cell count map. Factor weights were used to train a multiclass logistic regression classifier, which achieved an average classification accuracy of 59% (chance = 16.7%), indicating that semi-NMF effectively captured condition-dependent ensemble activity (**Figure 64B**).

**Figure 5.**
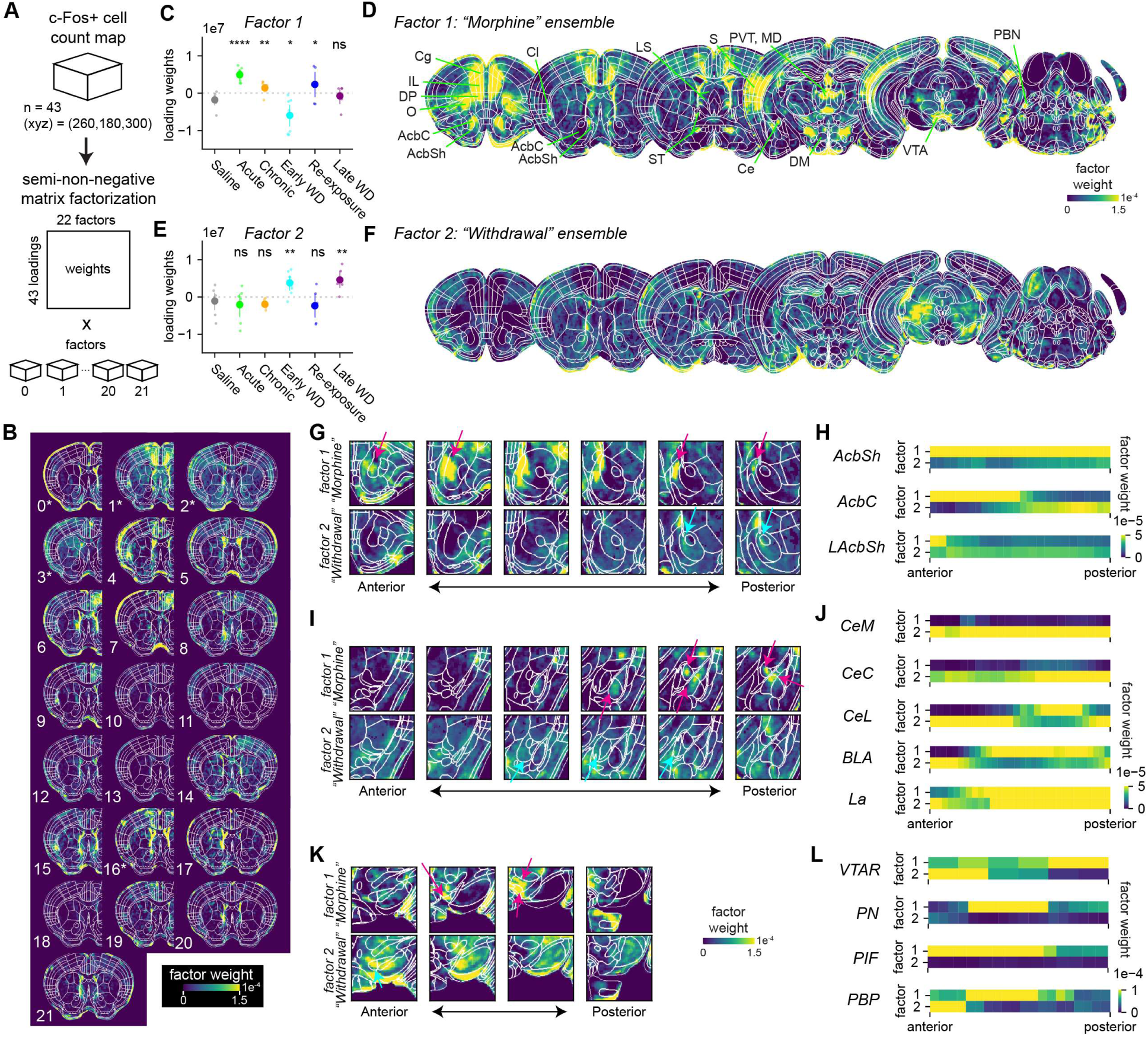
Voxel-wise factorization analysis reveals spatially distinct morphine administration and withdrawal ensembles. (A) Schematic showing the semi-nonnegative matrix factorization analysis for the c-Fos+ cell density map. A total of 22 factors were identified. (B) Representative coronal section showing weights for 22 factors. Factor 0, 1, 2, 16 had significant changes in loadings compared to the saline group. Additional statistics are shown in **Table 5**. (C**–**F) Scatterplot showing loading weights for factor 1 (C) and factor 2 (E). Representative 50-μm coronal planes showing factor weights for factor 1 (D) and factor (F). (G**–** L) Representative 50-μm coronal planes showing factor weights for factor 1 and factor 2 in the nucleus of accumbens region (G), amygdalar region (I) and the ventral tegmental area (K) along the anterior-posterior axis. Heatmap showing average factor weights in the nucleus of accumbens region (H), amygdalar region (J) and the ventral tegmental area (L). Saline: n = 8, Acute: n = 7, Chronic: n = 8, EarlyWD: n = 8, Re-exposure: n = 6, LateWD: n = 6.

To further identify drug-related factors, we performed a one-way ANOVA followed by *post-hoc* comparisons of loading weights between saline and morphine-related drug conditions (**Figure S6C**). We found that factors 0,1, 2, 3 and 16 had significant changes in weights, suggesting that these factors represent neural activity associated with drug conditions. Interestingly, Factor 1 and Factor 2 showed contrasting weight distributions. In factor 1, we found significantly increased weights in acute, chronic and re-exposure conditions but significant decrease in earlyWD condition (**Figure 5C** and **5D**). In contrast, in factor 2, we found a significant increase in earlyWD and lateWD conditions (**Figure 5E** and **5F**). These results suggest that factors 1 and 2 capture a general morphine-administration and withdrawal ensemble, respectively. Interestingly, we found that the spatial pattern of these factors aligned with the “hotspot–coldspot” model^8–11^ of hedonic value coding. In the accumbens, which is shown to contain hotspots in the anterior AcbSh and coldspots in posterior AcbSh^12,14^, we found that signals from factor 1 were distributed from the anterior part of AcbC and AcbSh to the posterior part of AcbSh (**Figure 5G** and **5H**). In contrast, signals from factor 2 were absent in the AcbSh but concentrated in the posterior part of the AcbC. In the ventral pallidum (VeP), factor 1 matched the posterior hedonic hotspot, whereas factor 2 also engaged this region, raising the possibility that a coldspot-like territory coexists within the same anatomical domain (**Figure S7A and S7B**). In the insular cortex, previously reported posterior hedonic hotspots^39^ were replaced by a more global increased factor 1 signal across granular (GI), dysgranular (DI), and agranular (AI) subregions during morphine administration, with minimal increase in factor 2 (**Figure S7C and S7D**). Orbitofrontal cortex (OFC) signals similarly exceeded predicted hotspot boundaries^39^, spanning medial, ventral, and lateral OFC in factor 1(**Figure S7E and S7F**). In the amygdala, while the region-based analysis indicated significant increase in early withdrawal conditions (**Figure 4E and 4F**), our analysis revealed a concentrated signal in the posterior central amygdala lateral nucleus (CeL), a region implicated in suppressing aversive responses^20,34^, as well as medial subdivisions of the lateral (La) and basolateral (BLA) nuclei (**Figure 5I and 5J**). Factor 2 was broadly found across the anterior-posterior axis in Ce, La, BLA, importantly with a reduction in the posterior CeL and medial La and BLA where the factor 1 signal was found. In the VTA, factor 1 signals were localized in the medial to posterior part of the paranigral nucleus (PN), parainfudibular nucleus (PIF) and parabrachial pigmented nucleus (PBP) subregions while factor 2 were localized more anterior in the VTAR subregion (**Figure 5K and 5L**). In contrast, we found regions such as claustrum (Cl) and paraventricular thalamic nucleus (PVT) that are primarily associated with morphine-administration, to have a wide spread of signal in factor 1, but no signal in factor 2 (**Figure S7G–J**). Together, these results show that semi-NMF resolves spatially precise and functionally distinct ensembles for morphine-administration and withdrawal, revealing state-specific organization within classic reward and aversion circuits at subregional resolution.

### Morphine administration and withdrawal ensembles are distinct at the cellular level

While the unbiased spatial classification revealed distinct regional patterns of morphine-related neural activation, this approach alone cannot resolve whether the underlying neural ensembles overlap at the cellular level. To address this, we performed brain-wide, cellular-resolution activity mapping using the targeted recombination in active populations^40^ (TRAP) system, which enables permanent genetic labeling of neurons with tdTomato which were active during a defined time window governed by tamoxifen administration (**Figure 6A**). Using TRAP along with c-Fos immunolabeling, we were able to label two neural ensembles in the same animal. Each subject went through two phases of drug administration, one to genetically label neural activity by TRAP (TRAP phase), and the other to molecularly label neural activity using c-Fos antibody (c-Fos phase). We generated four experimental groups: acute^TRAP^ (TRAP: acute + Antibody: acute), saline^TRAP^ (saline + acute), chronic^TRAP^ (acute + chronic), and earlyWD^TRAP^ (acute + early withdrawal) (**Figure 6B**). Brains were collected, cleared, and stained for c-Fos, then imaged using light-sheet microscopy to detect c-Fos+, tdTomato+, and double-labeled (double+) cells (**Figure 6C**). To assess the overlap of the two ensembles in Acute^TRAP^ and others, we used a GLM framework to compare double+ cell density across experimental conditions while adjusting for biological covariates (**Figure 6D**). For each brain region, we fit a full model including chronic^TRAP^, earlyWD^TRAP^ and saline^TRAP^ and covariates such as sex, body weight, and age. We then fit a reduced model excluding treatment conditions and used a likelihood ratio test (LRT) to determine whether inclusion of treatment improved model fit. The resulting p-values were input to the BRANCH, which identified significant discoveries distributed throughout the brain, and 90 significant regions among the 233 curated brain region list. We first evaluated TRAP labeling performance using two metrics: averaged accuracy (double+ / total c-Fos+ cells) and efficiency (double+ / total tdTomato+ cells) using data from the 90 significant brain regions. The acute^TRAP^ group showed significantly higher accuracy than saline^TRAP^ and earlyWD^TRAP^, but not chronic^TRAP^ (**Figure 6E**). Similarly, TRAP efficiency was significantly greater in acute^TRAP^ compared to earlyWD^TRAP^, but not the other groups (**Figure 6F**). To integrate these two measures into a single index, we calculated the TRAP fidelity score, defined as the harmonic means of accuracy and efficiency (Fidelity=2 × (efficiency + accuracy) / efficiency × accuracy). The acute^TRAP^ group showed significantly higher fidelity than saline^TRAP^ and earlyWD^TRAP^, indicating more specific and efficient labeling of morphine-activated ensembles (**Figure 6G**).

**Figure 6.**
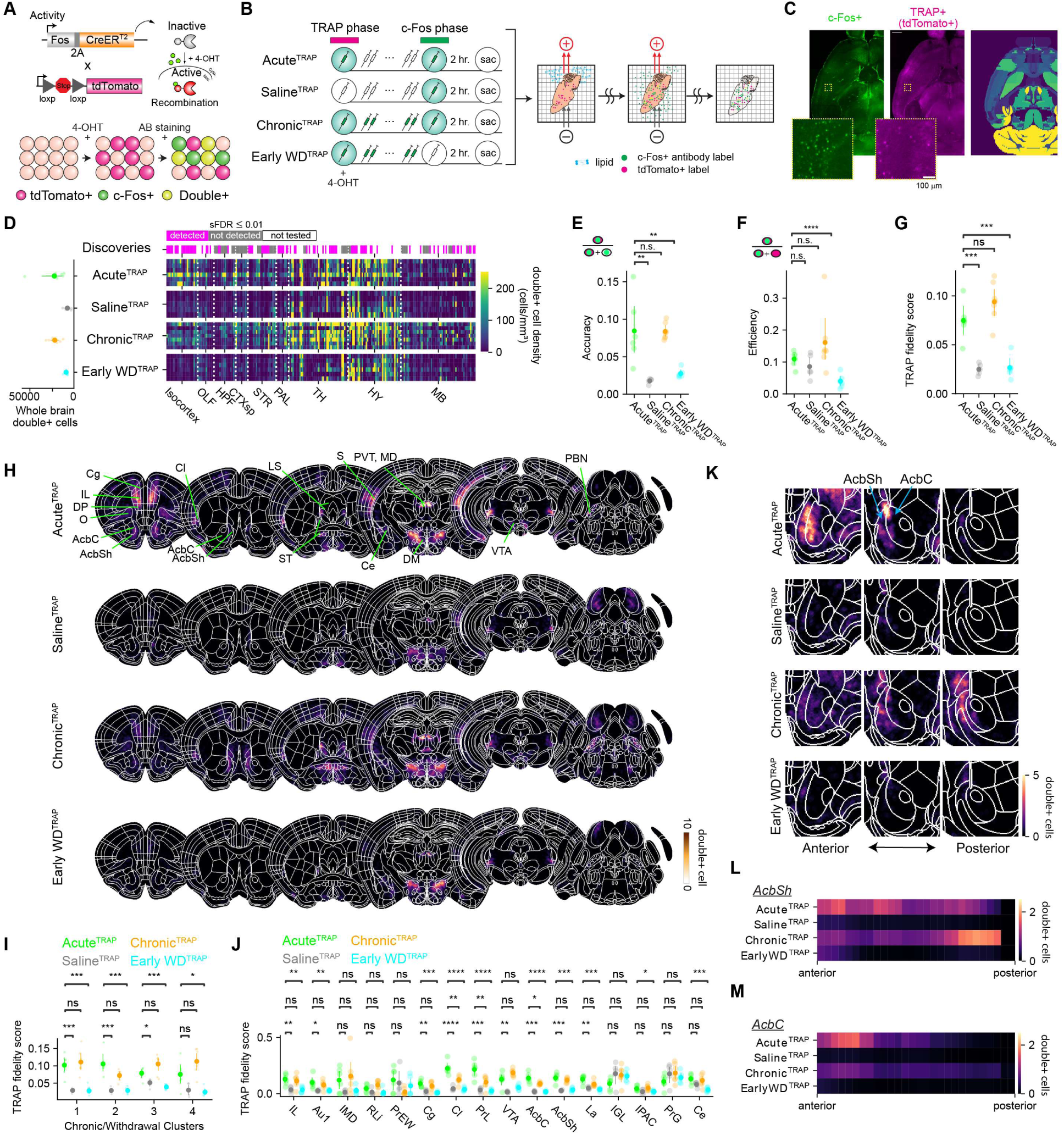
Analysis of acute morphine specific neural ensemble using TRAP2. (A) Schematic illustration showing genetic and molecular labeling of two neural ensembles within the same animal using TRAP2 and c-Fos+ antibody. Neurons that were active in both neural ensembles are expected to be tdTomato+ and c-Fos+ (double+). (B) Overview of the experiment pipeline. To examine the overlap of neural ensembles responding to acute morphine and to other drug conditions, we prepared 4 conditions which went to 4 different sets of drug administration schedules. Subjects went through two drug schedules, first for TRAP labeling and second for c-Fos immunolabeling. Acute^TRAP^: acute morphine ➔ acute morphine. Saline^TRAP^: saline ➔ acute morphine. Chronic^TRAP^: acute morphine ➔ chronic morphine. EarlyWD^TRAP^: acute morphine ➔ early withdrawal. Two hours after the second drug schedule, the subjects were perfused to collect brain tissue. Brain tissues were further processed to be cleared and stained, then imaged on a light sheet microscope. (C) c-Fos+ and tdTomato+ cells were registered to the Unified mouse brain atlas for further quantification. (D) Scatter plot showing total number of double+ cells in each subject (left) and heatmap showing double+ cell density for each brain region. (E**–**G) Scatter plot showing metrics of TRAP2; (E) accuracy (number of double+ cells / number of tdTomato+ cells). (F) efficiency (number of double+ cells / number of c-Fos+ cells), (G) TRAP fidelity score (Fidelity=2 × (efficiency + accuracy) / efficiency × accuracy). Students t-test with Benjamini-Hochberg correction. (E) Acute^TRAP^ vs. Saline^TRAP^: *P* = 7.59 × 10^-3^, *t*(9) = 3.42. vs. Chronic^TRAP^: *P* = 0.96, *t*(11) = 0.046. vs. EarlyWD^TRAP^: *P* = 9.44 × 10^-3^, *t*(9) = 3.20. (F) Acute^TRAP^ vs. Saline^TRAP^: *P* = 0.24, *t*(9) = 1.26. vs. Chronic^TRAP^: *P* = 0.23, *t*(11) = -1.27. vs. EarlyWD^TRAP^: *P* = 3.46 × 10^-4^, *t*(9) = 5.30. (G) Acute^TRAP^ vs. Saline^TRAP^: *P* = 3.22 × 10^-4^, *t*(9) = 5.63. vs. Chronic^TRAP^: *P* = 0.11, *t*(11) = -1.76. vs. EarlyWD^TRAP^: *P* = 3.08 × 10^-4^, *t*(9) = 5.39. (H) Representative 50-μm coronal planes showing average double+ cells. (I) Scatter plot showing average double+ cell density by subject among brain regions in each cluster identified in Figure 4. Student’s t-test was conducted between Acute^TRAP^ and every other condition with post-hoc Benjamini/Hochberg correction over conditions. Cluster 1: Acute^TRAP^ vs. Saline^TRAP^: *P* = 0.044, *t*(9) = 2.35. vs. Chronic^TRAP^: *P* = 0.10, *t*(11) = -1.80. vs. EarlyWD^TRAP^: *P* = 0.013, *t*(9) = 3.02. Cluster 2: Acute^TRAP^ vs. Saline^TRAP^: *P* = 5.42 × 10^-4^, *t*(9) = 5.23. vs. Chronic^TRAP^: *P* = 0.70, *t*(11) = 0.39. vs. EarlyWD^TRAP^: *P* = 3.80 × 10^-4^, *t*(9) = 5.24. Cluster 3: Acute^TRAP^ vs. Saline^TRAP^: *P* = 4.19 × 10^-3^, *t*(9) = 3.80. vs. Chronic^TRAP^: *P* = 0.30, *t*(11) = -1.09. vs. EarlyWD^TRAP^: *P* = 1.60 × 10^-3^, *t*(9) = 4.28. Cluster 4: Acute^TRAP^ vs. Saline^TRAP^: *P* = 1.97 × 10^-4^, *t*(9) = 6.02. vs. Chronic^TRAP^: *P* = 0.42, *t*(11) = 0.84. vs. EarlyWD^TRAP^: *P* = 2.72 × 10^-4^, *t*(9) = 5.47. Cluster 5: Acute^TRAP^ vs. Saline^TRAP^: *P* = 6.11 × 10^-4^, *t*(9) = 5.14. vs. Chronic^TRAP^: *P* = 0.044, *t*(11) = 2.27. vs. EarlyWD^TRAP^: *P* = 2.88 × 10^-4^, *t*(9) = 5.43. Cluster 6: Acute^TRAP^ vs. Saline^TRAP^: *P* = 6.82 × 10^-3^, *t*(9) = 3.49. vs. Chronic^TRAP^: *P* = 0.077, *t*(11) = -1.95. vs. EarlyWD^TRAP^: *P* = 2.01 × 10^-4^, *t*(9) = 5.69. (J) Scatter plot showing TRAP fidelity score in selected brain regions. Full results are shown in **Figure S8**. Student’s t-test was conducted between Acute^TRAP^ and every other condition with post-hoc Benjamini/Hochberg correction. Additional statistics are shown in **Table 6**. (K) Representative 50-μm coronal planes showing double+ cells in the accumbens region along the anterior-posterior axis. (L**–** M) Heatmap showing average double+ cell density in accumbens shell (L) and core (M) along the anterior-posterior axis. Acute^TRAP^: n = 6. Saline^TRAP^: n = 5. Chronic^TRAP^: n = 7. EarlyWD^TRAP^: n = 6.

We then examined the anatomical distribution of double+ cells. As expected, in acute^TRAP^ mice, the spatial pattern of TRAP-labeled neurons closely resembled the acute morphine activation map (**Figure 6H**). Notably, enrichment in prefrontal cortex, nucleus accumbens, medial thalamus, and hindbrain regions was also observed in the chronic^TRAP^ group, but not in saline^TRAP^ or earlyWD^TRAP^, suggesting that acute and chronic morphine largely recruit overlapping ensembles, while early withdrawal engages a distinct population. We next evaluated how the acute morphine administration ensemble captured by TRAP is represented by the functional clusters of brain regions identified in **Figure 4** (**Figure 6I**). These clusters reflected consistent spatial modules that were differentially recruited during chronic morphine exposure or early withdrawal. We found the TRAP fidelity score was significantly higher in acute^TRAP^ conditions than in the saline^TRAP^ and earlyWD^TRAP^ conditions, both in chronic morphine-dominant clusters (clusters 1, 2 and 4) and in earlyWD-dominant cluster (cluster 3). This result suggests that acute morphine administration cells are distinct from the withdrawal-responsive cells in most brain regions. Individual regional analysis further confirmed this (**Figure 6J** and **Figure S6**). Interestingly, we found that regions such as Cl and PrL had significantly larger TRAP fidelity score in the acute^TRAP^ condition than in the chronic^TRAP^ condition suggesting that the chronic exposure to morphine may attenuate neural activity in these specific brain regions. Spatial mapping along the anterior-posterior axis of the nucleus accumbens revealed that acute^TRAP^ and chronic^TRAP^ animals had broad recruitment in both AcbC and AcbSh, whereas earlyWD^TRAP^ showed minimal double⁺ labeling, demonstrating that morphine and withdrawal activate largely non-overlapping neuronal populations even within the same anatomical (**Figure 6K–L**). The acute^TRAP^ and chronic^TRAP^ groups displayed broad recruitment across AcbC and AcbSh. In contrast, earlyWD^TRAP^ animals showed minimal double+ cell labeling throughout the accumbens, suggesting that morphine administration and withdrawal engage largely distinct neuronal populations even within the same anatomical region. Together, these findings show that TRAP combined with c-Fos immunolabeling captures acute morphine administration neurons with high fidelity and reveals that morphine administration and withdrawal ensembles are distinct across the brain at cellular resolution.

### Molecular markers for morphine related neural ensembles in the accumbens

Whole-brain mapping of neural activity enabled us to identify spatial distribution of neural activity after morphine-administration or withdrawal. To identify molecular markers of neurons activated by these conditions, we performed a voxel-wise Pearson correlation analysis between spatial gene expression patterns from the Allen Brain Institute MERFISH dataset^41^ (2.8 million cells, 147 coronal sections, 1122 gene panel, 5322 cell-type clusters) and condition-specific predictive activation maps within a brain region of interest (**Figure 7A**). For each morphine-related condition, we computed correlations coefficient between gene expression levels and predictive activity maps, identifying genes whose spatial distributions significantly aligned with condition-specific neural activation (**Figure 7B–J**; **Figure S9**). We conducted this correlation analysis in 3 key brain regions identified in the preceding analysis that contained condition specific neural ensemble patterns; the Acb, Ce and VTA. In the Acb, among the genes with significant correlation, we identified genes such as *Kcnip1* and *Stard5* as markers for the acute-ensemble in the AcbSh, *Neurod3* as chronic-ensemble in the anterior Acb and *Calcr* as a marker for the earlyWD marker in the posterior AcbC (**Figure 7B–D; Figure S9A**). Importantly, this correlation coefficient was larger than classical cell type markers in the Acb such as *Drd1* and *Drd2*. In the Ce, we identified *C1ql2* and *Prkcd,* a classically known cell type marker of the CeL which regulates fear expression^42^, to correlate with the localized signal found in the acute-ensemble (**Figure 7E–G; Figure S9B**). Other CeA markers such as *Calcrl* or *Crh* had weaker correlation with this signal. We additionally found *Otx2* and *Prlr* as potential markers of global Ce activation during the earlyWD state. For the VTA, *Calb1, Chrna6* and *Th,* a cell type marker for dopaminergic cells in the VTA, showed expression patterns that significantly correlate with the acute morphine ensemble (**Figure 7H–J; Figure S9C**). This correlation coefficient was larger than that of excitatory and inhibitory cell type markers, *Slc17a6* and *Slc32a1. Npas1* was found for anterior VTA but not posterior VTA cell type markers.

**Figure 7.**
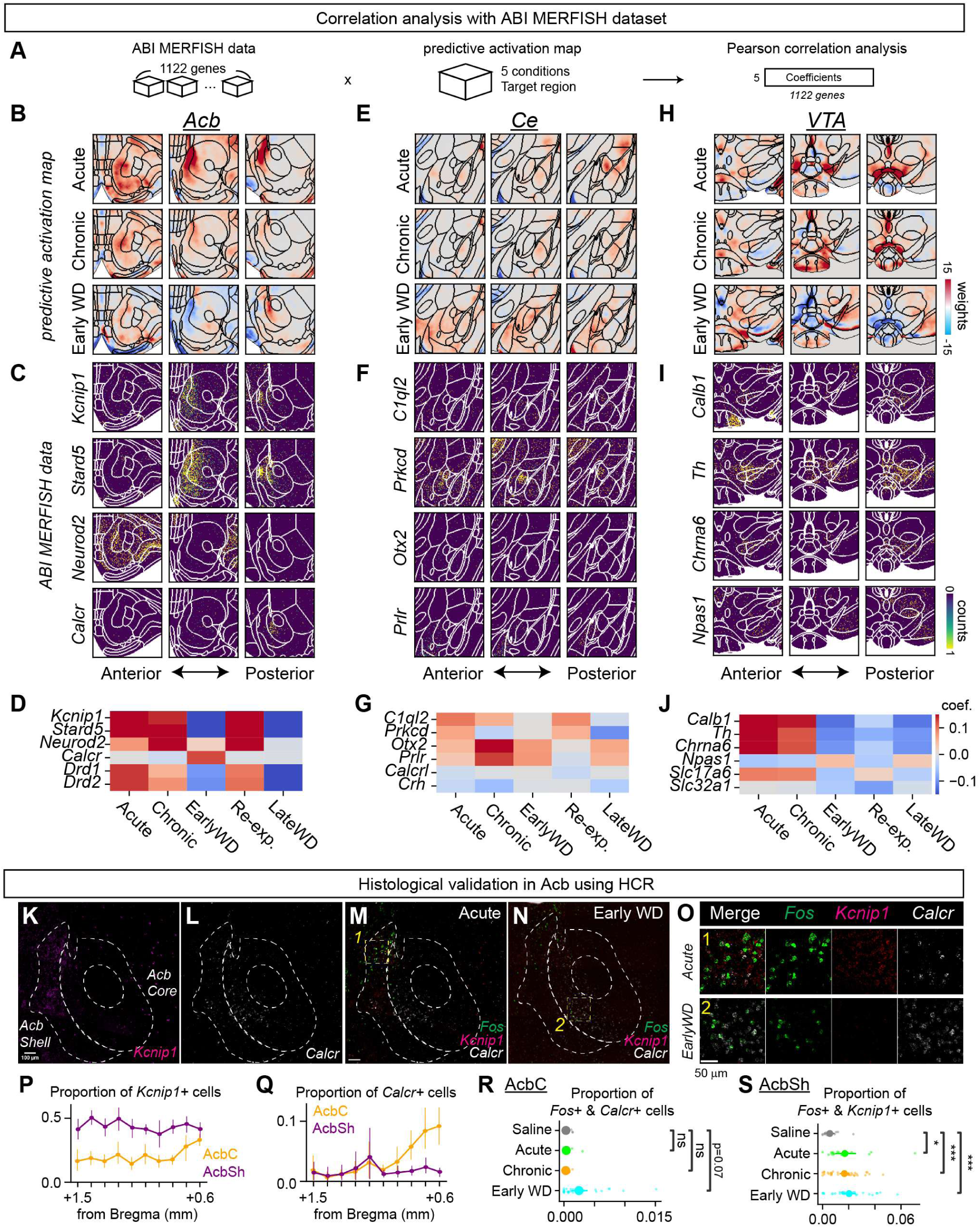
Identification of key molecular markers using c-Fos+ predictive activity map. (A) Schematic illustration of gene expression correlation analysis. Pearson correlation coefficients were calculated using c-Fos predictive activity map in the accumbens for each experimental condition and spatial gene expression data extracted from the ABI MERFISH dataset. (B**–**J) Results from the correlation analysis. Representative 50-μm coronal planes showing c-Fos predictive activity map in acute, chronic and early withdrawal conditions and 6 representative molecular markers in the (B and C) nucleus accumbens (Acb), (E and F) central amygdala (Ce) and (H and I) ventral tegmental area (VTA). Heatmap showing coefficient value for selected genes In (D) Acb, (G) Ce and (J) VTA. List of 20 genes with highest correlation coefficients are shown in **Figure S9**. (K**–**L) Representative images showing HCR *in situ* staining of 20-μm coronal sections with the Acb labeling (K) *Kcnip1* and (L) *Calcr*. Scale bar, 100 μm. (M–N) Representative images showing *Fos, Kcnip1* and *Calcr* expressing cells in the Acb in (N) acute morphine administration and (N) early withdrawal conditions. Scale bar, 100 μm. (O) Enlarged area of Acb highlighted in yellow in J and K. Scale bar, 50 μm. (P**–**Q) Line plot showing distribution of the proportion of *Kcnip1*+ cells (P) and *Calcr*+ cells (Q) in the accumbens (+1.5 mm to +0.6 mm from Bregma). Orange line, AcbC. Purple line, AcbSh. (R**–**S) Strip plot showing (R) proportion of *Calcr+* and *Fos+* cells in the AcbC (Saline vs. Acute: *P* = 0.83, *t*(31) = -0.21. vs. Chronic: *P* = 0.84, *t*(53) = 0.20. vs. EarlyWD: *P* = 2.32 × 10^-2^, *t*(44) = -2.35.) and (S) proportion of *Kcnip1+* and *Fos*+ cells in AcbSh (Saline vs. Acute: *P* = 1.63 × 10^-2^, *t*(31) = -2.54. vs. Chronic: *P* = 4.75 × 10^-4^, *t*(53) = -3.73. vs. EarlyWD: *P* = 2.85 × 10^-4^, *t*(44) = -3.94.). Saline: n = 16 sections, Acute: n = 17 sections, Chronic: n = 39 sections, EarlyWD: n = 30 sections.

To further evaluate the transcriptional identity of neurons activated by morphine-administration or withdrawal, we conducted a linear regression analysis using the cell-type cell count map derived from the Allen Brain Institute MERFISH dataset^41^ and whole-brain cell-type atlas^43^. The Allen mouse whole-brain cell-type atlas transcriptionally and spatially identifies 5322 clusters as their finest resolution of cell-type. Predictive activity maps for the Acb, Ce and VTA in each morphine-condition were modeled as a linear combination of cell-type cell count maps (**Figure S10A; Figure S11 and S12**). In Acb, we first found that a subcluster of dopamine receptor 1-expressing cells (0964 STR D1 Gaba_9) significantly contributed to the acute, re-exposure and chronic morphine administration neural activity signal (**Figure S10B**). This cluster is spatially located in the medial part of the AcbC and AcbSh, medial part along the AP-axis (**Figure S10C and S10D**) and uniquely expresses *Tac1* and *Pdyn* as well as *Drd3* as molecular markers. Another cluster with significant contribution was a cluster located in the olfactory tubercle and the medial part of AcbSh and uniquely expresses *Drd3* and *Stac* (0939 OT D3 Folh1 Gaba_1). In the chronic morphine model, we found 0964 STR D1 Gaba_9 cluster, as well as a glutamatergic cluster which labels cells in the anterior pat of the Acb (0171 IT AON-TT-DP Glut_2). In the early withdrawal model, we found one cluster from both dopamine receptor 1 and 2 -expressing cell-types (0962 STR D1 Gaba_8 and 0985 STR D2 Gaba_5) that were localized to the posterior part of the AcbC and are labeled by *Dio3* and *Calcr* respectively. Importantly, 0964 STR D1 Gaba_9 which was a significant cluster in the acute morphine administration model was found to have a significantly negative contribution in the early withdrawal model, strengthening our finding that the morphine-administration and withdrawal ensembles in the Acb are spatially, cellularly and molecularly distinct. In the Ce, we found that 1332 CEA-BST Six3 Cyp26b1 Gaba_2 cluster, localized to the CeL subregion and uniquely expresses *Prkcd*, significantly contributed to the acute morphine model (**Figure S11**). In the VTA, we found that multiple clusters of dopaminergic neurons (3849 SNc-VTA-RAmb Foxa1 Dopa_2, 3840 SNc-VTA-RAmb Foxa1 Dopa_1, 3852 SNc-VTA-RAmb Foxa1 Dopa_2, 3842 SNc-VTA-RAmb Foxa1 Dopa_1) significantly contributed to the acute morphine model (**Figure S12**).

Lastly, to validate this systematic approach to identify molecular identities of a given neural ensemble, we performed multiplexed hybridization chain reaction^44^ (HCR) to assess co-expression of immediate early gene *Fos* and the marker genes. We targeted the Acb, and examined *Fos*, *Kcnip1*, and *Calcr* co-expression in coronal brain sections collected 30 minutes after drug administration across acute, chronic, early withdrawal, and saline conditions **(Figure 7H–P**). Consistent with spatial transcriptomic predictions, *Kcnip1* expression spanned the AcbSh across the anterior-posterior axis (**Figure 7H and 7M**), while *Calcr* was localized to the posterior AcbC (**Figure 7I and 7N**). Co-labeling revealed that *Calcr+Fos+* cells were selectively enriched in the posterior core during early withdrawal, indicating that *Calcr* serves as a marker for withdrawal-responsive neurons ((**Figure 7J–L and 7P**). In contrast, *Kcnip1+Fos+* cells were found throughout the shell under acute, chronic, and early withdrawal conditions, suggesting that *Kcnip1* marks morphine administration responsive neurons as well as a subset of withdrawal-responsive cells in the AcbSh (**Figure 7J–L and 7O**). Together, these results highlight the utility of integrating whole-brain activity maps with spatial transcriptomics to molecularly define functional neural ensembles.

## Discussion

Understanding how opioids reshape brain function requires mapping neural activity across the entire brain with sufficient resolution to resolve the underlying cellular ensembles. Here, we combined whole-brain c-Fos mapping, a hierarchical statistical framework (BRANCH), and complementary molecular and genetic tools to define how morphine administration and withdrawal recruit distinct but spatially, cellularly, and molecular organized networks. This multiscale approach revealed that administration and withdrawal states engage largely non-overlapping neuronal populations, even within the same anatomical region, and that these ensembles can be further distinguished by spatial topography, molecular markers, and receptor expression profiles. Our findings not only clarify the large-scale organization of opioid-related brain states but also provide a generalizable pipeline for hierarchically and spatially dissecting complex behavioral conditions.

This comprehensive whole-brain activity dataset spanning multiple morphine-related conditions with cellular resolution allowed us to link brain-wide ensemble engagement with molecular identity. To systematically detect condition-related changes, we applied BRANCH, a hierarchical statistical testing approach that accounts for the brain’s nested anatomical organization. The BRANCH testing approach was able to improve sensitivity without sacrificing interpretability, enabling the detection of convergent subthreshold signals across related subregions that would be missed by conventional statistical testing. Although applied here to c-Fos activity maps, this framework could be readily extended to other whole-brain datasets, including projection mapping, cell-type–specific activity recordings, and whole-brain transcriptomics.

### Region-specific suppression of MOR+ cells and the logic of disinhibition

Despite the widespread expression of mu-opioid receptors (MOR) across the brain^7^, morphine-induced neural activation is strikingly selective. In contrast to the brain-wide expression of MOR (**Figure S8**), the distribution of the hedonic opioid hotspot-coldspots^12–14^ as well as the neural activity evoked by acute morphine administration were found in concentrated brain regions (**Figure 2**). This suggests that morphine induces brain wide activity through a multi-layered regulation model, where receptor presence is a prerequisite, but how the receptors are expressed within a local circuit architecture shape actual recruitment. Furthermore, since MOR is a Gi-coupled receptor, this local circuit architecture requires MOR to be expressed in inhibitory neurons to induce disinhibition of a larger neural ensemble. In our data, by using whole brain labeling of MOR+ and KOR+ neurons, we find that the likelihood of MOR+ cells to be activated by morphine is generally like that of KOR+ neurons, suggesting that most brain regions do not contain this specific local circuit architecture (**Figure 3**). Interestingly, 3 out of 40 brain regions tested had a significant decrease in the likelihood of MOR+ cells being activated, in other words, a MOR+ neuron specific suppression of neural activity was only observed in 3 out of 40 brain regions: VTA, DR and IL. As proposed in previous studies, VTA is considered to have the local circuit architecture where MOR+ is biased to inhibitory neurons within the VTA^20,28–31^ or in the rostromedial tegmental nucleus^29^ (RMTg) which results in disinhibition dopaminergic output neurons by opioids, with the exception of a small subpopulation of DA neurons which show inhibition through MORs. Indeed, our molecular analysis revealed that the spatial pattern of acute morphine administration ensembles strongly correlates with the expression pattern of *Th*, a molecular marker of DA neurons, supporting the idea which morphine disinhibit DA neurons (**Figure 7**). These results further suggest that regions strongly activated by acute morphine, such as the Acb, is likely to be a secondary effect from the VTA local circuit activation. In the DR, MOR expression is observed in both excitatory and inhibitory cells^45,46^, but it is also reported that MORs disinhibit serotonin efflux^47–50^ suggesting that a local circuit architecture similar to the VTA may exist in the DR. This circuit structure is not identified in the IL, however, it is considered that MORs are expressed in non-pyramidal neurons^7^ in the PFC, suggesting that a disinhibitory circuit may exist. Nevertheless, additional studies are required to evaluate whether MORs in VTA, DR or IL regulate the local and the brain-wide neural ensemble evoked by morphine. Future studies could use highly receptor specific agonists such as fentanyl or DAMGO to induce MOR+ specific intervention or to employ brain region- and cell type-specific loss of function of MOR+ during morphine administration to identify the sufficiency or necessity of the receptors in these regions for the global neural ensemble. It is also important to note that our approach cannot detect opioid actions at the pre-post synaptic terminal which is known to have an important role in neural modulation^51^. Future studies using whole brain antibody labeling of MOR/KORs could examine the correlation of the spatial localization of these receptors and brain wide neural ensembles. Additionally, extending this receptor-based labeling approach to withdrawal conditions, which is suggested to evoke aversive states via KOR+ neural ensembles^52–55^, could reveal whether the recruitment logic of these ensembles shifts across drug states, potentially illuminating distinct cellular mechanisms that underlie the transition from reward to aversion. Overall, our results emphasize the utility of hierarchical and spatial whole-brain activity mapping pipeline as a powerful framework to systematically study the cellular logic of drug action.

### Revisiting the hotspot–coldspot model in a brain-wide framework

Understanding the brain-wide neural responses to different opioid-related conditions has been critical for developing effective interventions for OUDs. Whole brain mapping of opioid-related neural activity has partially been conducted in several studies, but they lacked comprehensive analysis of the spatial and cellular distribution of opioid administration and withdrawal ensembles^17–22^. Here, we applied region level analysis with BRANCH-based correction and voxel level regression and semi-NMF analysis to conduct detailed investigation of the hierarchical and spatial distribution of morphine and withdrawal associated neuronal ensembles. Importantly, our analysis provided an opportunity to evaluate the opioid hotspot–coldspot model in a systematic fashion. We decomposed c-Fos+ maps into latent spatial factors that captured ensemble structure (**Figure 5**). Morphine-related ensembles (e.g., factor 1) and withdrawal-related ensembles (e.g., factor 2) were spatially segregated aligning with the hotspot–coldspot hypothesis^12–14^. Indeed, our study allowed us to revisit this hypothesis with fine detail in the mouse brain which led to novel insights. While classically the AcbSh has been extensively studied as the hotspot subregion, in the mouse brain morphine-related ensembles are not only encoded in the AcbSh but also in the medial part of the AcbC. Our data demonstrates that the withdrawal-related ensemble is mainly located in the posterior AcbC, while the studies in rat brain found the opioid hedonic coldspots to be in the AcbSh. These fine characterizations of the spatial map of potential hotspots-coldspots demonstrate the strength of the systematic review of neural ensembles. Other regions which were suggested to contain opioid hedonic hotspot-coldspots, such as the medial orbitofrontal cortex (OFC) and posterior insula^39^, showed inconsistent results in our study (**Figure S5**). For example, the insular cortex exhibited broad activation with morphine but lacked clear topology. There are several possible explanations for these discrepancies. First, c-Fos is an integrative marker of neural activity with limited temporal resolution and may not distinguish neural responses related to the hedonic value of morphine from those reflecting interoceptive, cognitive, or homeostatic processes. As such, regions like the insula and OFC, known to multiplex value, sensory, and emotional signals, may exhibit composite patterns of activity not resolved by c-Fos labeling. Second, our analysis detects increases in activity; however, opioids are potent suppressors of neuronal firing via MOR activation. Thus, functionally important reductions in neural activity, especially within MOR-expressing neurons, may be underrepresented or missed entirely. This raises the possibility that certain coldspot regions may encode opioid-related signals through suppression rather than excitation. Together, these considerations highlight both the power and the limitations of c-Fos-based whole-brain activity mapping and suggest that complementary tools such as neural inhibitory indicators such as pPDH^56^ are needed to further characterize the opioid hedonic hotspot-coldspot landscape.

Additionally, while the extensive studies in rats regarding the opioid hedonic hotspot-coldspots have provided important understandings to the encoding of valence, the field has lacked functional intervention studies of this model, mainly due to the lack of means to conduct detailed molecular characterization in rats. In contrast, our whole mouse brain dataset allows integration of multi-modal datasets to further characterize the properties of the opioid hedonic hotspot-coldspots. As an example, by aligning the *in situ* transcriptional data provided by Allen brain institute^41,43^ with our neural ensemble data, we demonstrated a powerful *in silico* pipeline to identify molecular identities of morphine-administration and withdrawal ensembles. We focused on the Acb, Ce and VTA and first identified genes that have correlated expression level with morphine-related neural ensembles (**Figure 7**). We then expand this analysis by utilizing the transcriptionally and spatially defined cell-type information provided through the Allen whole-brain cell-type atlas^43^. This analysis allowed us to identify cell-type clusters that are likely to contribute to the morphine-related ensembles. For example, in the context of the opioid hedonic hotspot-coldspot in the Acb, our analysis first found collection of genes such as *Kcnip1* and *Calcr* to be significantly correlated with the morphine- administration and withdrawal ensembles respectively (**Figure 7B and 7C**). We then identified that the ensembles consist mainly of 0964 STR D1 Gaba_9 cluster for the administration ensemble and 0962 STR D1 Gaba_8 and 0985 STR D2 Gaba_5 clusters for the withdrawal cluster (**Figure S10**). Indeed, data from the Allen brain cell atlas^43^ shows that 0964 STR D1 Gaba_9 cluster shows high levels of *Kcnip1* expression, while 0985 STR D2 Gaba_5 cluster specifically shows high level expression of *Calcr* compared to other Acb clusters. We further validate this finding using *in situ* hybridization analysis (**Figure 7H–P**). These molecular markers should serve as a key genetic handle to characterize the sufficiency and necessity of the opioid administration and withdrawal-related responses. Overall, these results highlight the strength of brain-wide mapping of neural activity to a standardized brain atlas allowing integration of multimodal datasets collected by the neuroscience community for detailed characterization of neural ensembles. However, it is also important to note that this approach requires the target neural ensembles to localize at the voxel-level and fails to detect sparse signals that are spatially intermingled with other cell-types or other neural ensembles. Additional methodological advancements to increase the spatial resolution of analysis and to conduct robust and multiplexed labeling approaches for mRNA and proteins in the whole-brain level are equally important to create a truly comprehensive *in silico* analysis pipeline.

Taken together, our study defines a distributed and modular organization of brain-wide neural ensembles that underlie distinct phases of opioid experience. By bridging hierarchical and spatial scales, we show that morphine and withdrawal engage fundamentally separate neural ensembles. This work provides a foundation for understanding how the brain encodes the opposing motivational forces of reward and aversion and offers a generalizable strategy for decoding the logic of complex internal states at the whole-brain level.

## Materials and methods

### Animals

All experiments were approved by the Institutional Animals Care and Use Committee at the University of Washington (Protocol #4450-01). Animals were group-housed with littermates on a 12-hr light cycle at ∼22°C with food and water available *ad libitum*. Wild-type C57BL/6 mice were purchased from the Jackson laboratory and bred in-house. Ai14 (Rosa-CAG-LSL-tdTomato-WPRE, Jax#007914) were purchased from Jackson laboratory. TRAP2 mice^57^ (also known as Fos-2A-iCreER^T2^) were provided by Dr. Palmiter (University of Washington) and crossed with Ai14 to generate TRAP2::Ai14 mice. MOR-Cre (also known as Oprm1Cre:GFP knock-in/knock-out, Jax#035574) mice were provided by Dr. Palmiter. KOR-Cre^58^ (also known as Oprk1-Cre, Jax#035045) mice were provided by Dr. Bruchas (University of Washington).

### Drug administration

We used pharmaceutical-grade morphine procured from UW drug services. Mice in chronic, re-exposure, early withdrawal and late withdrawal conditions were given an escalating dose of morphine over the course of 7 days (0.1, 0.3, 0.5, 1, 5, 10, 20 mg/kg). As for control, mice in saline and acute conditions were given saline injections. Each mouse received two i.p. injections at each dose ∼10-14 hours apart, for a total of 14 injections. A final dosage was given 10-14 hours after the last injection to the saline, acute, chronic and early withdrawal conditions and 21 days after the last injection to the re-exposure and late withdrawal conditions. For the final dosage, saline, early withdrawal and late withdrawal received saline. Acute, chronic and re-exposure received 20 mg/kg of morphine. Subjects were perfused for brain collection 2 hours after the final dosage.

### Retro-orbital labeling of MOR/KOR+ cells

AAV2/PHP.eB Ef1a-DIO-H2B-tdtomato-WPREs (BrainVTA, BHV17100164, #lot EB-6182-K240702, 1-2×10^13^ vg/ml) was diluted to 0.5×10^12^ vg/ml with PBS for a final volume of 100 μl and drawn up into a ½ ml 27-gauge tuberculin syringe. The animal was anesthetized with isoflurane in an induction chamber until unresponsive to toe pinch. The animal was then placed in left lateral recumbency so that the injector can protrude the right eyeball by pulling the dorsal skin and ventral to the globe from the socket with gentle pressure. The needle was then inserted at a 30–45-degree angle into the medial canthus to target the retro-bulbar sinus, bevel down to decrease likelihood of damaging the eye. The virus was injected within 30 seconds and then the needle was slowly withdrawn to minimize any bleeding. After the injection, the animal was placed back in its home cage for recovery. More than three-weeks after the injection, animals were then administered to drugs as described in **Drug administration.**

### TRAP experiments

TRAP method was conducted as previously described with modifications (**Figure 5** and **Figure S4**) ^57,59^. Each subject went through two phases of drug administration, one to genetically label neural activity by TRAP, and the other to molecularly label neural activity using c-Fos antibody. For the TRAP phase, each subject was habituated to injection 1-week prior to the TRAP day. 24 hours prior to the TRAP day, subjects were isolated in a new cage. On the TRAP day, the subjects were administered with the 1st drug during the first 3 hours of the dark cycle. 1-hour after the administration, the subjects were I.P. injected with 0.125 mL of 50 mg/kg of 4-Hydroxytamoxifen (Sigma-Aldrich, cat#H6278) dissolved in sesame oil. After the injection, the subjects were kept in their isolated cage. After 24 hours once the tamoxifen activity is reduced, the subjects were returned to their co-housing cages. Immediately after, the animals followed their drug schedule for the antibody labeling phase. Seven days after the TRAP day, the subjects were isolated in a new cage. On the antibody labeling day, the subjects were administered with the 2nd drug. 2-hours after the administration, the subjects were anesthetized to perfuse for brain extraction.

### HCR experiments and analysis

Hybridization chain reaction (HCR) was performed based on the protocol provided by Molecular Instruments with some modifications (Figure 7E–L). To induce *Fos* labeling, subjects were isolated in a new cage overnight. On the following day, the subjects were administered with the 2nd drug. 30 minutes after the administration, the subjects were immediately sacrificed to collect brain tissue. The brain tissue was then snap-frozen using dry ice. Every fifth 20-μm coronal sections containing the Acb were collected on a slide glass and fixed with 4% PFA in PBS for 30 min and rinsed with PBS, followed by dehydration with a series of 50%, 70%, 100%, 100% of ethanol. After the final dehydration the slides were dried. The tissue was treated with Proteinase K (10 U/mL, Biolabs, P8107S, lot#10140231) for 5–7 minutes. The slides were then rinsed with PBS and were applied with hybridization buffer (Molecular Instruments) for 10 minutes at 37°C for pre-hybridization. RNA probes for Round1: *Fos, Oxtr* and Round2: *Kcnip1, Calcr* (1 mM, Molecular Instruments) were diluted in a ratio of 1:250 into the hybridization buffer. After pre-hybridization, hybridization buffer (70 μl) containing the probes was applied to each slide on which the parafilm cover was placed. After 12h–18h of incubation at 37°C in the moisture chamber, the sections were washed, with a series of probe wash buffer and SSC-T buffer (5x SSC, 0.1% Tween 20) mixture. After the final wash, the sections were incubated in amplification buffer (Molecular Instruments) for 30 minutes at room temperature for pre-amplification. Amplification hairpin probes (3 mM, Molecular Instruments) were diluted in a ratio of 1:50 into the hybridization and then snap cooled for 1.5 minutes in 95°C. After pre-amplification, amplification buffer containing hairpin probes (70 μl) was applied to each slide on which the parafilm cover was placed. After 12h–18h of incubation at room temperature in the moisture chamber, the sections were washed in 5xSSC-T buffer for 30 minutes x2. The sections were quenched for autofluorescence using Vector TrueView reagent (Vector, cat# SP-8400). After washed in 2xSSC, the sections were stained with DAPI in PBS (5 μg/ml, Thermo Fisher Scientific, cat#D1306) for 8 minutes. The slide was then mounted using Vibrance Antifade Mounting Medium (VECTASHIELD, cat#H-1800). Images were obtained using a conventional microscope (Zeiss, Apotome Imager.M2) with a 20x objective. After image acquisition, the coverslip was removed and washed in 2x SSC for 10 minutes, then incubated with DNase1 (0.25 U/μl, Roche, cat#4716728001) for 1.5 h at room temperature. The section was then washed in 2x SSC for 5 minutes x6. The sections then proceeded to the pre-hybridization step for the next round of HCR. A total of 2 rounds of HCR were conducted.

Images from multiple rounds were aligned with a custom Python script using opencv2 findHomography. Cells were segmented based on the DAPI signal using cellpose package and their pretrained cyto2 model^60^. mRNA signals were detected using big-FISH package with spot radius set to 500 nm × 500 nm^61^. Parameters for detecting puncta and intensity of each gene were manually adjusted for each tissue section of each animal. The cutoff value to determine a gene “expressing cell” was defined as 5 transcripts.

### Whole-brain clearing, active antibody staining and imaging

For brain wide c-Fos activity mapping, we adapted tissue clearing and active antibody staining methods provided from LifeCanvas SmartBatch+ system. Animals were prelabeled with fluorophores for the TRAP2 experiments (Figure 5) or KOR/MOR experiments (Figure 7), otherwise, a C57BL/6J mice were used. For brain collection, the animals were perfused with PBS and 4% PFA-PBS, the brain was harvested and post-fixed with PFA for 24-hours in 4°C with shaking. After post-fixation, the brains were processed for tissue clearing. For tissue clearing, we utilized stabilization under harsh conditions via intramolecular epoxide linkages to prevent degradation method (SHIELD)^23^. The procedures followed the SHIELD tissue clearing protocol (Full Active Pipeline Protocol v5.06, LifeCanvas). Briefly, the brain was incubated in SHIELD OFF solution [2.5 mL DI water, 2.5 mL SHIELD-Buffer Solution, 5 mL SHIELD-Epoxy Solution] at 4°C for 4 days. Then transferred to SHIELD ON buffer and incubated at 37°C for 24 hrs. Following, SHIELD ON buffer incubation, the brain was pre-incubated in Delipidization buffer for overnight at room temperature. The tissues were then placed into SmartBatch+ (LifeCanvas) to conduct active tissue clearing for 30-36 hours. After clearing, tissues were transferred to blocking buffer solution [200 μL normal donkey serum per brain, 0.5 mM Methyl-beta-cyclodextrin, 0.2% trans-1-Acetyl-4-hydroxy-L-proline in PBS + 0.02% sodium azide] for 48-72 hours at 37°C with shaking. Then, the tissues were pre-incubated in primary staining buffer (LifeCanvas) for 24 hours at RT. Following pre-incubation, the tissues were placed into SmartBatch+ (LifeCanvas) to conduct primary active tissue staining for 18 hours in primary staining buffer with 200 μL normal donkey serum and 1.75 ug of c-Fos antibody (Abcam, ab236039, lot#1073423) per brain. After staining, tissues were washed with PBS + 0.02% sodium azide for 8 hours, followed by overnight fixation with 4% PFA-PBS. For the secondary antibody staining, the tissues were incubated with secondary staining buffer for 8 hours at 37°C with shaking. Then the tissues were placed into SmartBatch+ to conduct secondary active tissue staining for 12 hours in secondary staining buffer with 200 μL normal donkey serum and 2.625 ug of Donkey Anti-Rabbit – SeTau-647 (LifeCanvas, DkxRb-ST647, lot#0324L500) per brain. After staining, tissues were actively washed with secondary staining buffer for 6 hours. Additionally, tissues were washed with PBS + 0.02% sodium azide for 4-8 hours, followed by overnight fixation with 4% PFA-PBS. To match the refraction index, the tissue was incubated in 50% of EASYINDEX solution (RI = 1.52, LifeCanvas) at 37°C shaking for 24 hrs and then 100% of EASYINDEX solution at 37°C shaking for 24 hrs. The tissue was then embedded in EASYINDEX with 2% agar and mounted onto SmartSPIM light sheet microscope (LifeCanvas). The tissue was mounted on the ventral side facing toward the objective, imaged from the horizontal view using a 1.6x objective with 4-μm zstep. 594 nm channel for tdTomato signals and 647 nm channel for c-Fos signals were collected, 488 nm channel was collected as the background channel. Background channel was used for the alignment to the brain atlas.

### Preprocessing of whole-brain datasets using high-performance computer clusters

Whole-brain images were first destriped and stitched using a custom script developed by LifeCanvas. Subsequently, images were registered to the Unified Mouse Brain Atlas^24^ at a resolution of 20 × 20 × 50 μm (x, y, z). c-Fos+ cells or tdTomato+ cells were segmented through the same pipeline using a built-in SpotDetection function or by using a classifier created using Ilastik ^62^. The cell coordinates together and meta information (cell signal intensity, size), were stored as a NumPy arrays. To train the classifier, 10 chunks of images were randomly sampled and used as the training dataset (Image size, (x,y,z) = 2.0 mm × 2.0 mm × 100 μm). One chunk image of the same size from each subject was used as the test dataset. The quality of the classifier was visually examined. The cell coordinated information in raw image space was projected to brain atlas space using the Elastic transformation matrix calculated during the registration process. Using the cell coordinate information in atlas space and the Unified Mouse Brain Atlas annotation, the number of cells as well as the density of the cells were calculated for each brain region. Cerebellum and hindbrain regions were omitted from the analysis. Using the cell coordinates in atlas space, we also calculated a voxelized cell count heatmap. By-voxel cell counts heatmap data and the by-region cell count, and cell density data were further used for downstream statistical analysis.

To identify and quantify spatial overlap between labeled cells from two imaging channels, we developed a custom Python script using scipy.spatial.cKDTree for efficient 3D spatial matching. Coordinates of c-Fos cells and tdTomato+ cells in raw image space were extracted as described above. Overlapping cells were defined based on proximity within a user-defined spatial tolerance (offset), which accounts for potential misalignments or biological variation. Specifically, for each cell in one array, the nearest neighbor in the second array was queried using a k-d tree structure, with overlap determined if the Euclidean distance between cells fell below the threshold defined by the offset parameter (8 × 8 × 4 μm).

The above preprocessing were done using a modified version of the ClearMap 1.0 pipeline ^24,63^ (available at: https://github.com/kenjp1223/ClearMap). This modification was designed to efficiently parallelize the preprocessing of mouse brain data across compute nodes on a high-performance computing (HPC) cluster. While ClearMap offers a robust pipeline for whole-brain analysis, it presents challenges in scalability and reproducibility. The original version requires certain software and hardware configurations and lacks native support for parallelization. To address these limitations, we provide a Singularity container bundling the modified ClearMap along with all necessary dependencies, ensuring streamlined installation and reproducibility across computing environments. Our version is specifically optimized to distribute preprocessing tasks for individual brain samples across multiple HPC nodes, enabling highly parallelized and scalable data processing. This approach significantly accelerates data throughput, particularly for large datasets. Moreover, using HPC infrastructure is often more cost-effective than upgrading local workstations, as it allows flexible scaling of computational cores, memory, and storage resources as needed. In this study, we leveraged compute resources provided by the Bridges-2 supercomputing cluster at the Pittsburgh Supercomputing Center via the National Science Foundation (NSF) ACCESS program^63–65^. The NSF ACCESS program offers substantial free computing credits for academic users, making it an ideal platform for whole-brain image processing. Together, our modified ClearMap pipeline and the HPC infrastructure available through NSF ACCESS offer a scalable, reproducible, and cost-efficient solution for preprocessing large-scale whole-brain imaging datasets.

### Generalized linear model and likelihood ratio testing

To assess the effect of experimental condition on c-Fos+ cell density across annotated brain regions, we fit generalized linear models (GLMs) using the statsmodels Python package. Each region was tested independently. The outcome variable was c-Fos+ cell density, and the primary predictor was experimental condition, modeled as a categorical variable with five levels: Acute, Chronic, Re-exposure, Early Withdrawal, and Late Withdrawal (Saline was treated as reference and excluded to avoid collinearity). Model specification:

Full model (Model A):

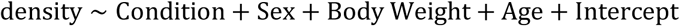

Reduced model (Model B):

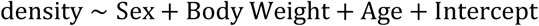

Both models were fit using Negative Binomial distribution to accommodate overdispersion in the count-derived response. Continuous covariates (Body Weight, Age) were scaled by mean. A likelihood ratio test (LRT) was performed by comparing the log-likelihoods of the two models:

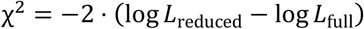

The p-value was computed using the chi-square distribution with degrees of freedom equal to the difference in the number of model parameters. Regions with insufficient subject counts or poor model convergence were excluded or assigned NaN.

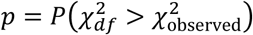

All statistical results (p-values per region) were compiled into a dataframe and subsequently passed into the BRANCH hierarchical testing pipeline for structured multiple hypothesis correction.

### Semi-nonnegative Matrix Factorization of c-Fos counts

To identify interpretable spatial patterns of brain-wide activity across animals, we applied a semi-nonnegative matrix factorization (semi-NMF) a previously developed framework^38^ to voxel-wise c-Fos cell count maps.

Briefly, the c-Fos+ cell count map was first downsampled to an isotropic 50 μm resolution. The input data consisted of non-negative integer cell counts *Y* ∈ *N*^M×D^, where *M* is the number of animals and *D* is the number of brain voxels (*M* = 43, *D* = 3932613). We utilized the algorithm described *Y*_m,d_ ∼ *poisson*(λ_m,d_) where the log-mean rates λ_m,d_ were parameterized through a generalized linear model using the nonlinear softplus function to ensure non-negativity:

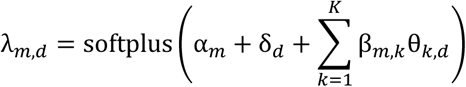

Here, *a*_m_ and δ_d_ are per-animal and per-voxel effects, respectively, β_m,k_ ∈ *R* are animal-specific loading weights for each of *K* shared latent factors θ_k_ ∈ Δ^D^. To improve stability, each update step was safeguarded using an Armijo backtracking line search. Initialization was performed using a non-negative singular value decomposition (NNSVD) on a variance-stabilized transformation of the count data. To determine whether individual semi-NMF factors were modulated by experimental condition, we performed a one-way ANOVA for each factor’s loading weights across all groups. Specifically, for each factor, we compared weights across saline and drug-exposed animals (acute, chronic, early withdrawal, re-exposure, and late withdrawal). ANOVA p-values were adjusted for multiple comparisons using the Benjamini–Hochberg false discovery rate (FDR) procedure across all 22 factors. For factors with significant FDR-corrected p-values (FDR < 0.05), we conducted pairwise post hoc Student’s t-test comparing each drug condition to saline.

### BRANCH hierarchical testing pipeline

In addition to mapping every voxel to a brain region, framworks for mouse brain atlases such as the Allen Brain Common Coordinate Framework^66^ or Unified Mouse Brain Atlas^24^ provide a hierarchical ontology of varying levels of refinement. This can be understood as a tree structure, where regions are nodes. The entire brain is the root node, coarse structures (e.g., gray matter) are children of the root node, and there are progressively finer resolved regions down the tree until we arrive at leaf nodes (i.e., regions that have no further subdivisions). To exploit this tree structure, we applied methodology proposed in Bogomolov. *et al*^25^. In brief, we (i) form a hierarchy of null hypotheses, (ii) associate p-values with all leaf nodes, (iii) propagate p-values up the tree using combination tests, and (iv) use the TreeBH method to test nodes *top-down* while controlling *sFDR*^l^ (defined below). We describe each of these steps in more detail below.

#### I: Tree Construction

Formally, we first associate a null hypothesis with each leaf node that its mean expression of c-Fos is invariant to treatment condition. We then form null hypotheses at coarser levels of resolution as the intersection of its child nulls (i.e., the null hypothesis associated with each non-leaf node is that *none* of its subregions have differential c-Fos expression due to treatment), and we repeat this process hierarchically up the tree.

#### II: Computation of leaf p-values

For each leaf node, we compute the per-mouse average c-Fos expression within that region. We fit a generalized linear model for each leaf region (as described above) and retain the p-value.

#### III: Propagation of p-values

The p-value associated with each non-leaf node is defined recursively: it is the result of applying the method of Simes^67^ to the p-values of all its child nodes. Given a sorted collection of *k* p-values *p*_(1)_ ≤ *p*_(2)_ ≤ ⋯ ≤ *p*_(K)_, Simes’ method returns a “combined” p-value as

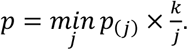

This combined p-value can be understood as corresponding to a test of the null hypothesis that *all* of the child null hypotheses are true.

#### IV: Testing with the TreeBH Method

With p-values attached to all of the nodes of the tree, we can proceed with the TreeBH method^25^. First, we set a targeted rate of control at each level of the tree (in practice, we set *q* = 0.01 at all levels). Starting with the root node, we apply the standard Benjamini-Hochberg method^68^ (BH) to the p-values of the child nodes at level *q*. We then proceed to the children of any rejected nodes and once again apply the BH method, but we use an adjusted *q* value based on the proportion of nodes under consideration (see Bogomolov et al.^25^ for more details). We continue recursively until we have no further rejections to pursue (or have reached leaf nodes).

#### Definition and Control of sFDR

Bogomolov et al. ^25^ introduced the *selective false discovery rate* at level l (*sFDR*^l^) in the context of hierarchical testing. While we provide a brief explanation of the key ideas here, we refer the reader to the source for a more thorough treatment. Mathematically, *sFDR*^l^ is defined as the expectation of the selective false discovery proportion (*sFDp*^l^). If we knew which of our discoveries were false, we could compute *sFDp*^l^ as follows. First, consider all rejected hypotheses at level l of the tree. Assign the value of 1 to those where the null was true (these are false discoveries) and assign 0 to those where the null was false (these are true discoveries). Next, at level l – 1, consider all hypotheses that were rejected. Assign to each the average of the values taken by its rejected children (if it has no rejected children, assign 0). Repeat this process for the l – 2 level and so on until we reach the root node: this value is *sFDp*^l^. While we cannot typically compute *sFDp*^l^ in practice (because we do not know which discoveries are true and which are false), it is nonetheless possible to demonstrate that under certain assumptions, the TreeBH algorithm will control the *expectation* of *sFDp*^l^, i.e., *sFDR*^l^, at a level no greater than a user-specified target *q*^l^, although in practice we choose a common value q to use at all levels and refer to *sFDR*^l^ throughout the manuscript simply as sFDR.

In Step III described above, we opted to propagate p-values using Simes’ method. While other methods such as Fisher’s can be more powerful, they are less robust to dependence, whereas Simes’ method remains valid under the forms of “positive” dependence likely to occur in imaging data as considered here. Moreover, the use of Simes’ method guarantees that TreeBH will give “consonant” results, i.e., that each non-leaf node that we reject will have at least one rejected child. Applied recursively, this guarantees that we will always be able to trace each rejected hypotheses to at least one leaf node. In addition, we are also able to invoke Theorem 1 of Bogomolov et al. ^25^, which implies control of *sFDR*^l^ for *all* levels. As a result, our error control holds even for “collapsed” trees, e.g., we can be confident of controlling sFDR at any level in the “sunburst” plots of **Figure 2**.

#### Software Implementation and Extension

While Bogomolov et al.^25^ offer a software implementation of their TreeBH algorithm, it is seemingly tailored to the setting wherein all leaf nodes occur at a common deepest level L. Because the hierarchical atlas commonly used in the neuroscience field does not satisfy this, we developed our own implementation that accommodates the native hierarchical structure of the mouse brain atlas. Moreover, internally it leverages tree data structures; this facilitates data import and visualization. The software is available on <u>GitHub</u> (https://github.com/Kessler-Friends/BRANCH).

### UMAP clustering analysis for chronic vs. early withdrawal

To identify patterns of brain-wide neural activity associated with chronic morphine and withdrawal conditions, we applied dimensionality reduction and clustering on by-region c-Fos+ cell count data. For each of the 233 curated brain regions, c-Fos+ cell densities were first extracted from 16 brains across experimental conditions (chronic morphine and early withdrawal). Density data was transformed using the natural logarithm to stabilize variance and mitigate the influence of high-weight outliers. Then the data were z-scored using sklearn.preprocessing. We then used Uniform Manifold Approximation and Projection (UMAP) to project the high-dimensional regional activity profiles into a 2-dimensional embedding space. UMAP was implemented using the umap-learn package (v0.5.3) with parameters: number of neighbors =3, minimum distance =0.2. To cluster brain regions based on their embedding in UMAP space, we applied hierarchical density-based spatial clustering of applications with noise (HDBSCAN, hdbscan v0.8.33), allowing the identification of distinct regional clusters that share similar morphine- or withdrawal-associated activation patterns.

### By-voxel analysis of c-Fos predictive activity mapping

To identify brain-wide spatial correlates of experimental condition, we performed voxel-wise linear regression using the statsmodels package in Python. This analysis was conducted at the level of individual voxels within the registered brain volume, leveraging spatial resolution to derive predictive maps of c-Fos expression.

At each voxel, the outcome variable was the c-Fos cell count across subjects (N_subject_ = 43, × = 650, y = 350, z = 300). This was further subset into voxels that contain brain data (N_voxel_ = 24393746). The predictors included experimental condition (Saline, Acute, Chronic, Re-exposure, Early Withdrawal, Late Withdrawal) and subject-level covariates (Sex, Body Weight, Age). Experimental conditions were modeled as a categorical variable with Saline as the reference group.

The regression model was specified as:

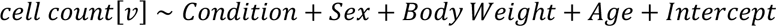

here *cell count*[*v*] is the c-Fos cell count at voxel v. All continuous covariates were mean centered before inclusion. The resulting voxel-wise regression weights (β coefficients) for each predictor were interpreted as c-Fos predictive activity maps, capturing the spatial distribution of each condition’s contribution to neural activation. These maps enabled unbiased examination of neural activity in order to asses which regions were most strongly modulated by morphine-related experimental conditions.

### Gene correlation analysis using Allen MERFISH data

To enable brain-wide voxel-wise comparison of gene expressions with other spatial datasets, we processed MERFISH data from the Allen Brain Institute’s Zhuang-ABCA-1 dataset (**Figure 7**) (Zhang et al., 2023), which provides single-cell resolution transcriptomic profiles from approximately 2.8 million cells across 147 coronal hemisphere sections throughout the mouse brain. This dataset includes log2-transformed gene expression counts, anatomical coordinates in the Allen Common Coordinate Framework (CCFv3, 25 µm resolution), and cell type annotations based on a hierarchical taxonomy. We accessed the data using the Allen Brain Cell Atlas API via the abc_atlas_access Python package. For each cell, we retrieved (1) log2-normalized gene expression values, (2) cluster assignments and metadata, and (3) anatomical coordinates in CCF space. To enable spatial alignment with our anatomical reference framework, we registered the Allen CCFv3 brain volume to the Unified mouse brain atlas (Kim et al., 2017; Chon et al., 2019), using symmetric normalization (SyN) image registration implemented in the Advanced Normalization Tools in Python (ANTsPy) framework. Specifically, we used a high-resolution Allen CCF coronal template as the fixed image and the Unified mouse brain reference as the moving image. The resulting forward transformation (comprising affine and nonlinear warp fields) was used for batch transformation of cellular coordinates. Single-cell anatomical coordinates were transformed from Allen CCF space to Unified mouse brain atlas space using ants.apply_transforms_to_points, taking into account voxel scaling (25 µm isotropic for CCFv3). Transformed coordinates were mapped to region annotations using the Unified mouse brain atlas label volume, allowing each cell to be associated with a discrete anatomical region in the Unified mouse brain atlas ontology. For each gene in the dataset, we extracted expression values and applied a positivity threshold defined as expression > 1. Cells passing this threshold were considered positive for that gene. To generate spatial expression volumes, we voxelized positive cells into a 3D image matching the Unified mouse brain atlas template dimensions (20 × 20 × 50 μm resolution). Each expressing cell contributed its expression value to surrounding voxels using a spherical kernel (40 × 40 × 100 μm voxels). This approach allowed for smooth spatial aggregation of discrete cell data into continuous voxel-level gene-expressing cell count map.

To identify the gene whose voxel-level gene-expressing cell count map correlates with morphine-induced neural activity, we conducted a voxel wise correlation analysis. First, voxel-level gene-expressing cell count maps for 1122 genes were restricted to hemisphere data. Similarly, the bilateral predictive activity map was averaged to generate a hemisphere predictive activity map. Then, the voxels were subset into ones that contained target region based on the Unified mouse brain atlas annotation. For each gene, we computed the Pearson correlation coefficient (r-value) and significance (p-value) between its voxel-level gene-expressing cell count map and the corresponding predictive activity map across voxels. P-values were corrected for multiple correction using Benjamini–Hochberg correction to identify significant correlation (alpha = 0.05).

### Cell type regression analysis using Allen MERFISH data

To enable brain-wide voxel-wise comparison of cell type distribution with other spatial datasets, we processed MERFISH data from the Allen Brain Institute’s Zhuang-ABCA-1 dataset (**Figure 7**) (Zhang et al., 2023) similarly as described in “**Gene correlation analysis using Allen MERFISH data**”. Briefly, using the transformed cell coordinate information and the cluster assignment data^43^, for each cell type, we voxelized positive cells into a 3D image matching the Unified mouse brain atlas template dimensions (20 × 20 × 50 μm resolution). Each expressing cell contributed its expression value to surrounding voxels using a spherical kernel (40 × 40 × 100 μm voxels). This approach allowed for smooth spatial aggregation of discrete cell data into continuous voxel-level by-cluster cell count map.

To identify clusters that predict morphine-induced neural activity, we conducted a voxel wise correlation analysis. The hemisphere predictive activity map was modeled as a linear combination of by-cluster cell count map using ordinary least squares regression with a constant term. By-cluster cell count map were first subset into ones that contained target region based on the Unified mouse brain atlas annotation. Then, these maps were further filtered to remove clusters with less than 10 cells. Predictors were z-scored column-wise, such that coefficients represent standardized effects (per 1 SD increase in cluster density). Robust (HC3) standard errors were used for inference, and cluster-wise significance was assessed by parametric *t*-tests with Benjamini–Hochberg correction (α = 0.05). To complement parametric inference, we performed coefficient-wise permutation testing by shuffling the target vector 500 times, recomputing regression coefficients, and estimating two-sided empirical p-values. Permutation p-values were also corrected using Benjamini–Hochberg (α = 0.05). Marker gene information was extracted from the allen brain cell atlas for additional characterization of cell-type clusters.

## Data distribution

Whole brain data is available on DANDI (https://dandiarchive.org/dandiset/001449/draft). All data used in this project is shared on Figshare (https://figshare.com/articles/dataset/Ishii-Morphine-2025/29497187). The code for analysis and data generation is shared on GitHub (https://github.com/stuberlab/Ishii-Morphine-2025).

## Author Contributions

Conceptualization, K.K.I. and G.D.S.; methodology, K.K.I., D.K. and G.D.S.; validation, K.K.I. and D.K.; formal analysis, K.K.I., Y.Z., L.R., S.B., A.H., S.C., H.P.S. and E.S.; coding, E.S. and A.D.L.; investigation, K.K.I., D.K., Z.C.Z., and E.R.S.; resources, S.A.G. and G.D.S..; data curation, K.K.I.; writing – original draft, K.K.I. and G.D.S.; writing – review and editing, K.K.I., D.K. and G.D.S. with input from all authors; visualization, K.K.I.; supervision, G.D.S.; and funding acquisition, G.D.S., D.K. and K.K.I.

## Acknowledgments

This work was supported by a JSPS oversea fellowship (K.K.I.), Uehara postdoctoral fellowship (K.K.I.), NIH grants R37 DA032750, R01 DA038168, R01 DA054317, P30 DA048736 and Washington Research Foundation Postdoctoral Fellowship (E.R.S.). D.K. was supported by NSF DMS-2303371, the Pacific Institute for the Mathematical Sciences, the Simons Foundation, and the eScience Institute at the University of Washington. E.S. was supported by NSF DMS-2134107. This work used Bridges-2 at Pittsburgh Supercomputing Center through allocation BIO230201, OTH240003, BIO240159 from the Advanced Cyberinfrastructure Coordination Ecosystem: Services & Support (ACCESS) program, which is supported by National Science Foundation grants #2138259, #2138286, #2138307, #2137603, and #2138296.

## Declaration of generative AI and AI-assisted technologies in the manuscript preparation process

During the preparation of this work the author(s) used ChatGPT and Claude for coding assistant, manuscript editing and proof reading. After using this tool/service, the author(s) reviewed and edited the content as needed and take(s) full responsibility for the content of the published article.

## Figures

**Figure S1.**
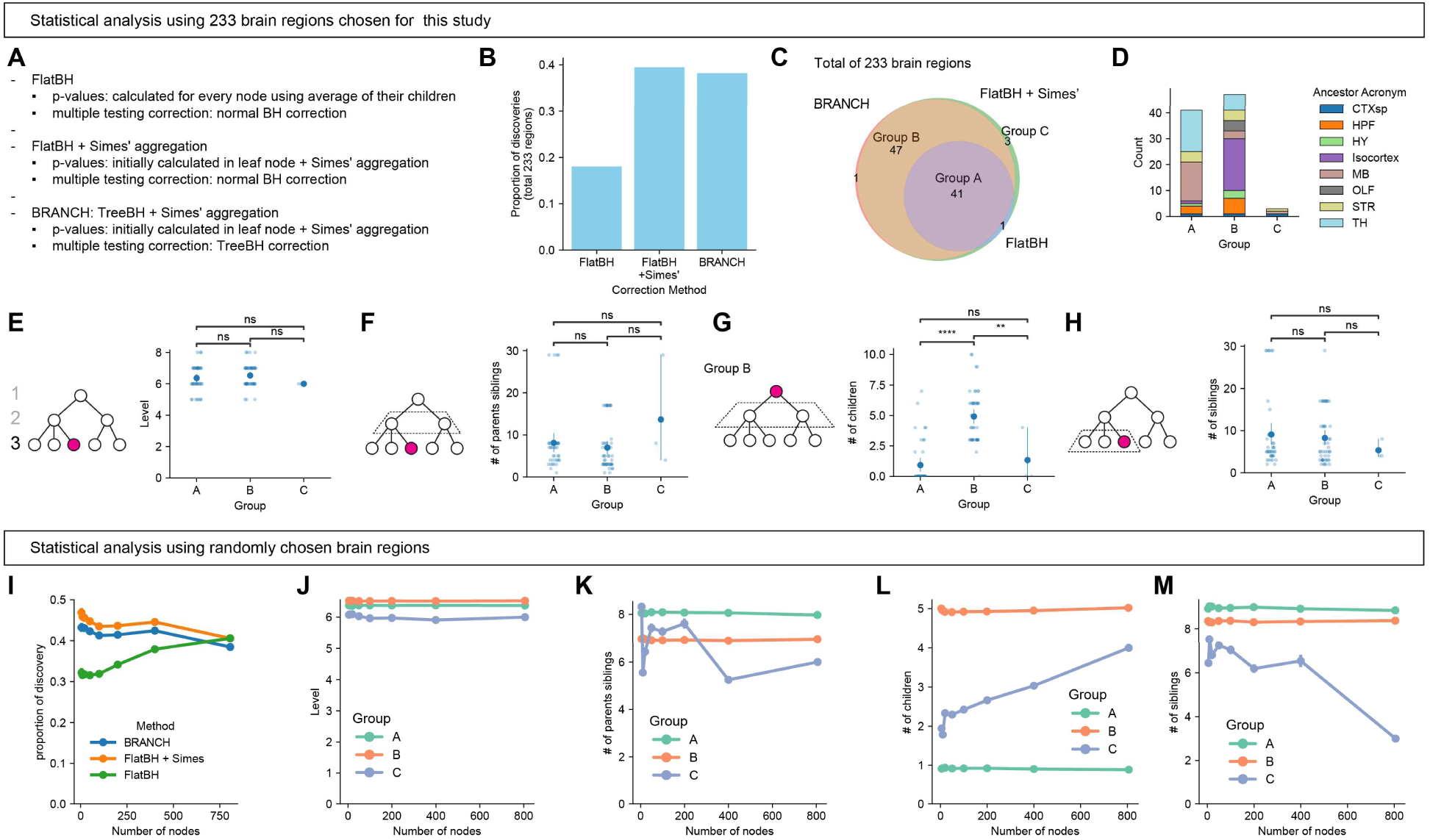
Comparison of BRANCH to classical statistical testing procedures, related to Figure 2. (A) Schematic overview of three procedures compared in this analysis. FlatBH: p-values were computed using average c-Fos+ cell density for each node. The list of p-values from the nodes of interest were processed with Benjamini–Hochberg (BH) correction. FlatBH + Simes: p-values were initially computed at leaf nodes, then were aggregated bottom-up using Simes’ aggregation method. The list of p-values from the nodes of interest were processed with BH correction. BRANCH: p-values were initially computed at leaf nodes, then were aggregated bottom-up using Simes’ aggregation method. P values from all the nodes were subjected to hierarchical sFDR correction using the TreeBH. (B) Bar graph showing the proportion of discoveries (out of 233 tested brain regions) made by each method. Both Simes’-aggregated methods (FlatBH + Simes and BRANCH) outperformed the non-Simes approach. (C) Venn diagram of statistically significant brain regions identified by each method. BRANCH and FlatBH + Simes’ shared a substantial number of discoveries not found by the FlatBH approach. Group A: Discovered by all three methods. Group B: Discovered by FlatBH + Simes and BRANCH, but not by FlatBH. Group C: Discovered by FlatBH + Simes only (i.e., not by BRANCH or FlatBH non-Simes). (D) Distribution of detected regions across anatomical divisions (ancestor acronyms) for each group of discoveries defined in (C). (E–H) Structural features of regions discovered by each method. (E) Depth in the hierarchy (levels from the root). (F) Number of sibling regions of the parent node (a proxy for branch width). (G) Number of children per discovered region. BRANCH and FlatBH + Simes preferentially identified parent nodes with many subregions, in contrast to FlatBH. (H) Number of sibling regions for each discovered node. Students t-test with Benjamini-Hochberg correction. (E) A vs. B: *P* = 0.33, *t*(90) = -0.97. B vs. C: *P* = 0.19, *t*(46) = 1.34. A vs. C: *P* = 0.39, *t*(50) = 0.86. (F) A vs. B: *P* = 0.49, *t*(90) = -0.69. B vs. C: *P* = 0.058, *t*(46) = -1.94. A vs. C: *P* = 0.19, *t*(50) = -1.33. (G) A vs. B: *P* = 1.7 × 10^-14^, *t*(90) = -0.91. B vs. C: *P* = 6.6 × 10^-3^, *t*(46) = 2.84. A vs. C: *P* = 0.46, *t*(50) = -0.74. Group A: n = 44, Group B: n = 48, Group C: n = 4. (I–M) Simulation results from analyses using randomly selected sets of brain regions (not based on biological data). (I) Proportion of discoveries as a function of the number of nodes sampled. (J–M) Structural properties of discovered regions across increasing tree sizes: depth (J), number of parent’s siblings (K), number of children (L), and number of siblings (M). Patterns confirm the generalizability of the effects observed in the curated 233-region set.

**Figure S2.**
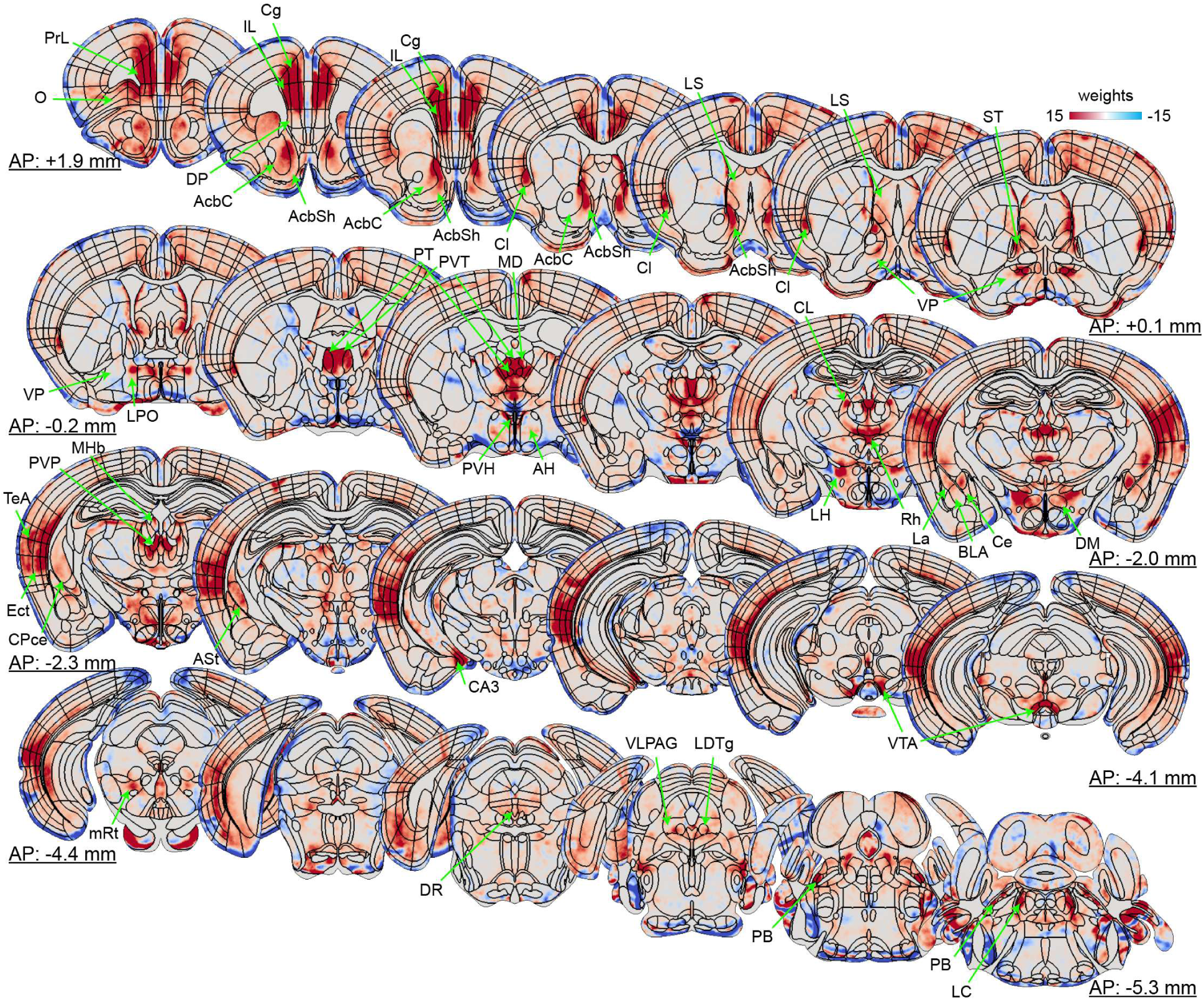
Neural activity pattern after acute morphine administration, related to Figure 3. Representative 50-m coronal planes showing predictive activity map for acute morphine condition.

**Figure S3.**
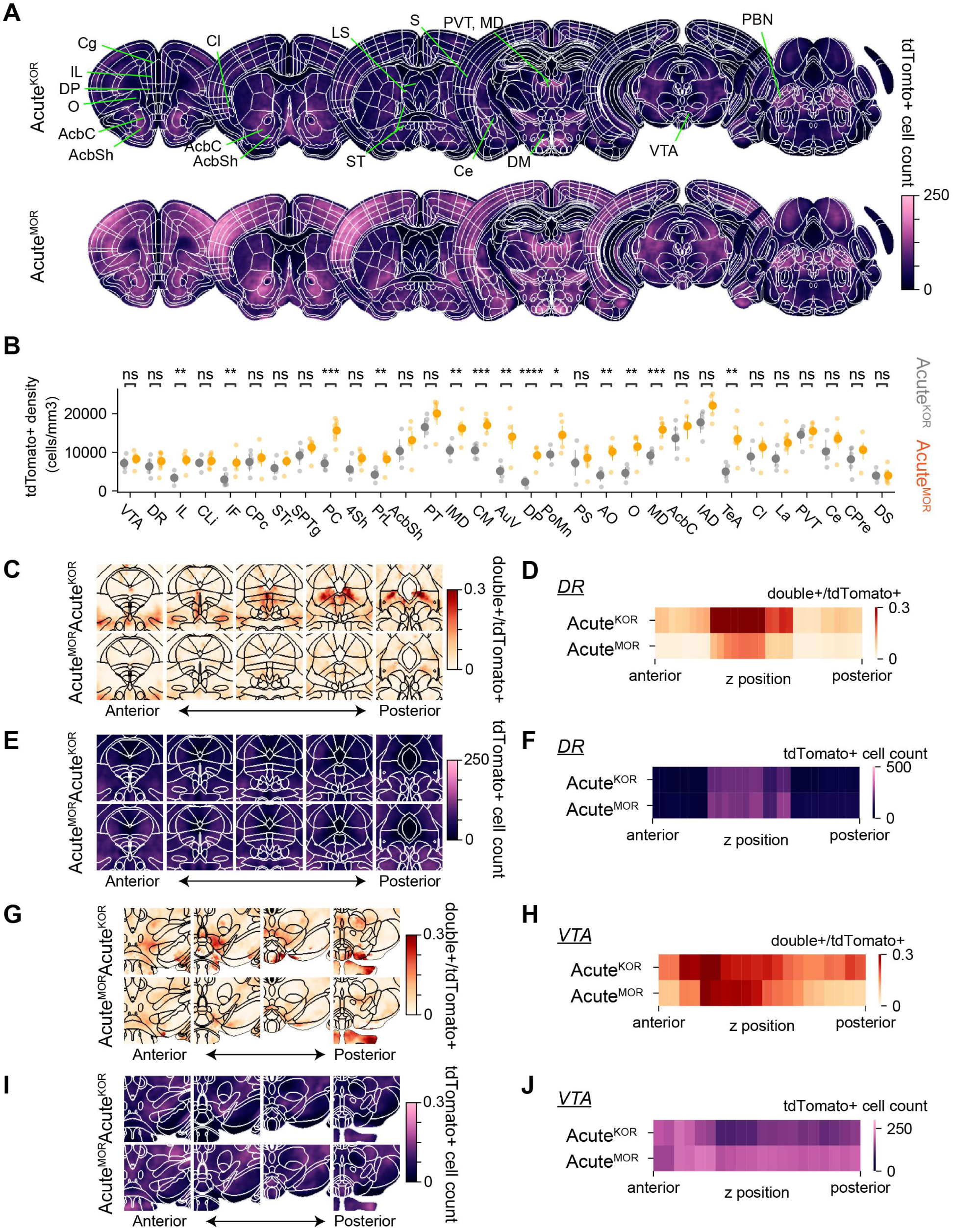
Characterization of opioid receptor expressing cells, related to Figure 3. (A) Representative 50-μm coronal planes showing average tdTomato+. (B) Scatter plot showing tdTomato+ among the 31 brain regions that were identified as acute morphine responsive regions in Figure 3B. Student’s t-test with post-hoc Benjamini/Hochberg correction over brain regions. (C) Representative 50-μm coronal planes showing double+ cells in the dorso raphe (DR) along the anterior-posterior axis. (D) Heatmap showing average double/tdTomato+ in DR. (E) Representative 50-μm coronal planes showing tdTomato + cells in the DR along the anterior-posterior axis. (F) Heatmap showing average tdTomato+ in DR. (G) Representative 50-μm coronal planes showing double+ cells in the ventral tegmental area (VTA) along the anterior-posterior axis. (H) Heatmap showing average double/tdTomato+ in VTA. (I) Representative 50-μm coronal planes showing tdTomato + cells in the VTA along the anterior-posterior axis. (J) Heatmap showing average tdTomato+ in VTA. Additional statistics are shown in **Table 4**.

**Figure S4.**
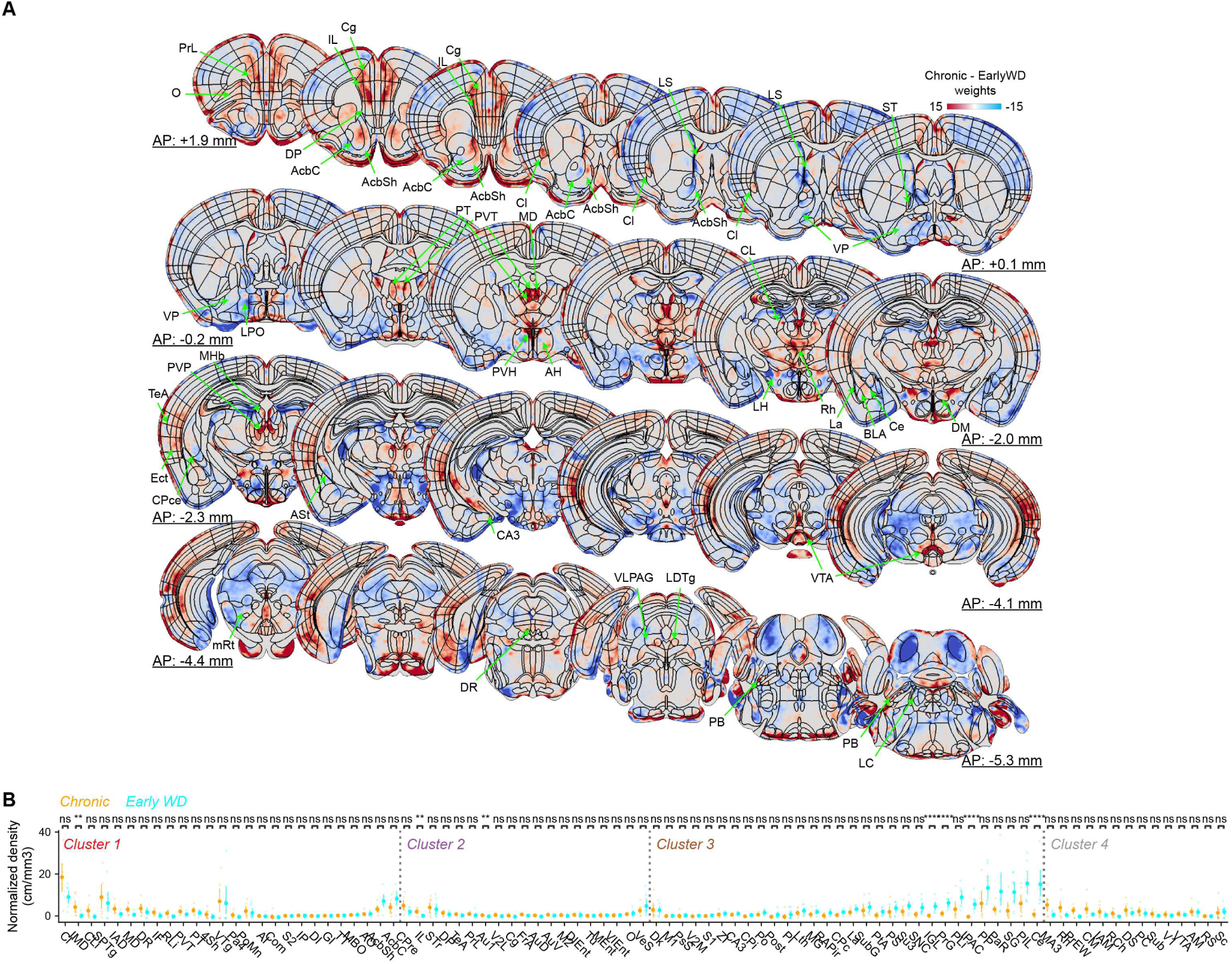
Distinct neural activity pattern during chronic morphine administration and early withdrawal, related to Figure 4. (A) Representative 50-μm coronal planes showing difference of predictive activity map between chronic and early withdrawal. (B) Scatter plot showing normalized c-Fos+ cell density in the chronic morphine and early withdrawal conditions among the 89 brain regions identified by BRANCH in Figure 2. Student’s t-test with Benjamini/Hochberg correction over brain regions (alpha = 0.05). Additional statistics are shown in **Table 3**.

**Figure S5.**
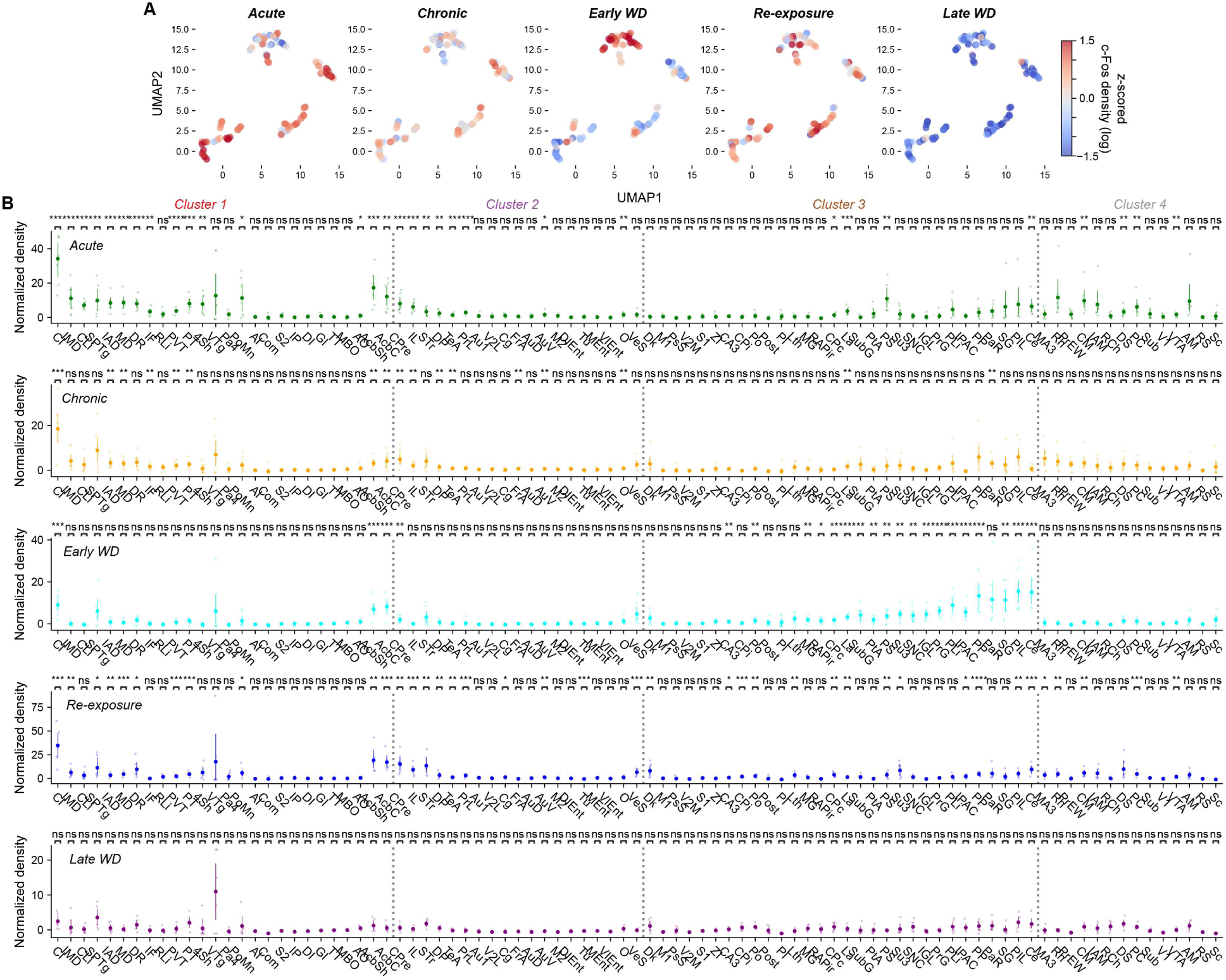
Normalized c-Fos+ cell density across brain region clusters, related to Figure 4. (A) UMAP display overlayed with z-scored c-Fos density. (B) Scatter plot showing c-Fos+ cell density normalized to saline in acute, chronic, early withdrawal, re-exposure and late withdrawal among the 89 brain regions identified by BRANCH in Figure 2. Student’s t-test with Benjamini/Hochberg correction over brain regions (alpha = 0.05). Additional statistics are shown in **Table 3**.

**Figure S6.**
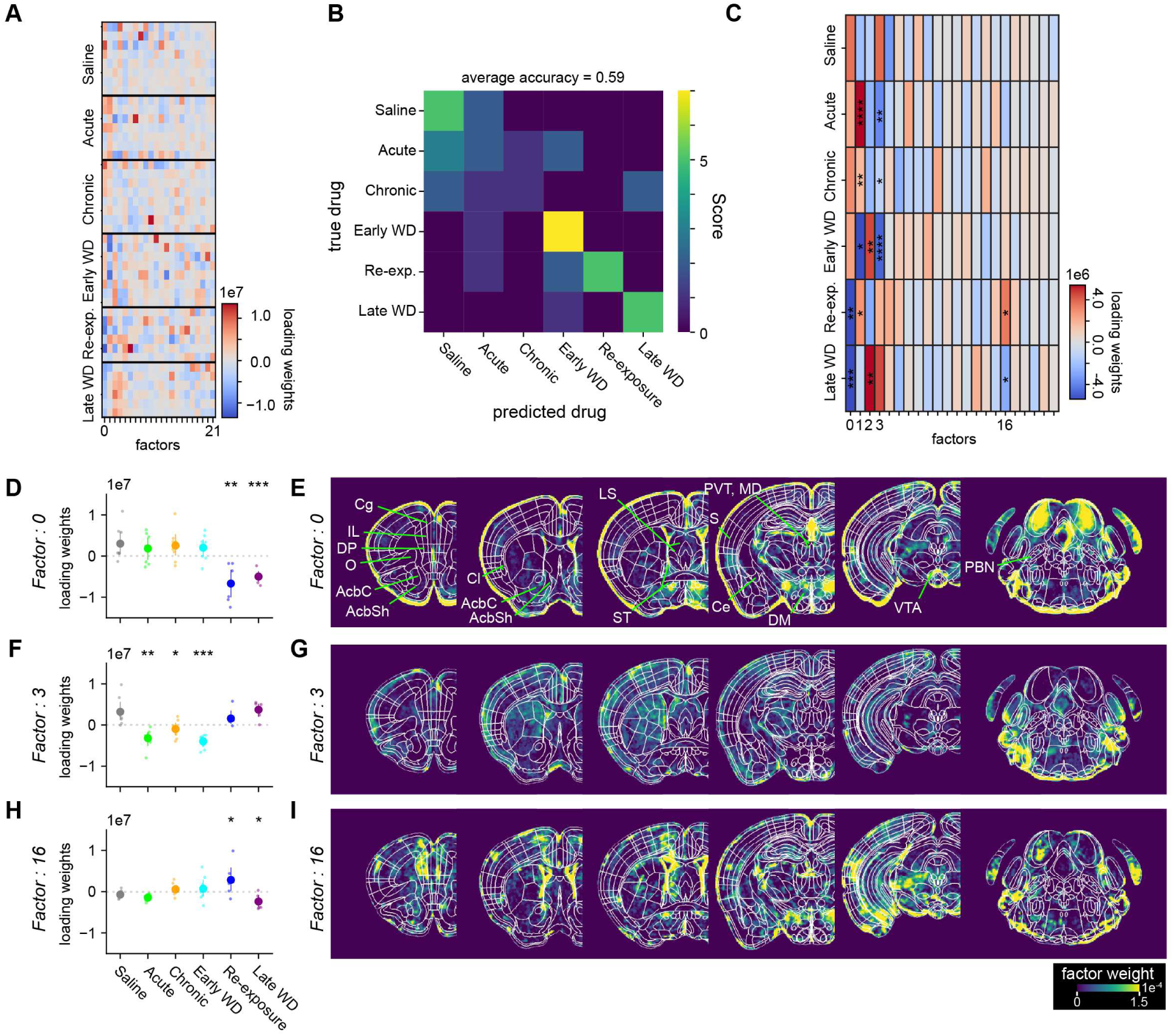
Voxel-wise factorization analysis to spatially classify distinct morphine responsive ensembles, related to Figure 5. (A) Heatmap showing loading weights for each drug condition in every subject. (B) Confusion matrix of drug condition classification based on loading weights. (C) Heatmap showing average loading weights for each drug condition in every factor. Factor 0, 1, 2, 16 had significant changes in loadings weights compared to the saline group. One-way ANOVA with post-hoc student’s t-test with Benjamini/Hochberg correction over conditions (alpha = 0.05). Additional statistics are shown in **Table 5**. (D–I) Scatterplot showing loading weights for factor 0 (D), factor 3 (F) and factor 16 (H) and representative 50-μm coronal planes showing factor weights for factor 0 (E), factor 3 (G) and factor 16 (I).

**Figure S7.**
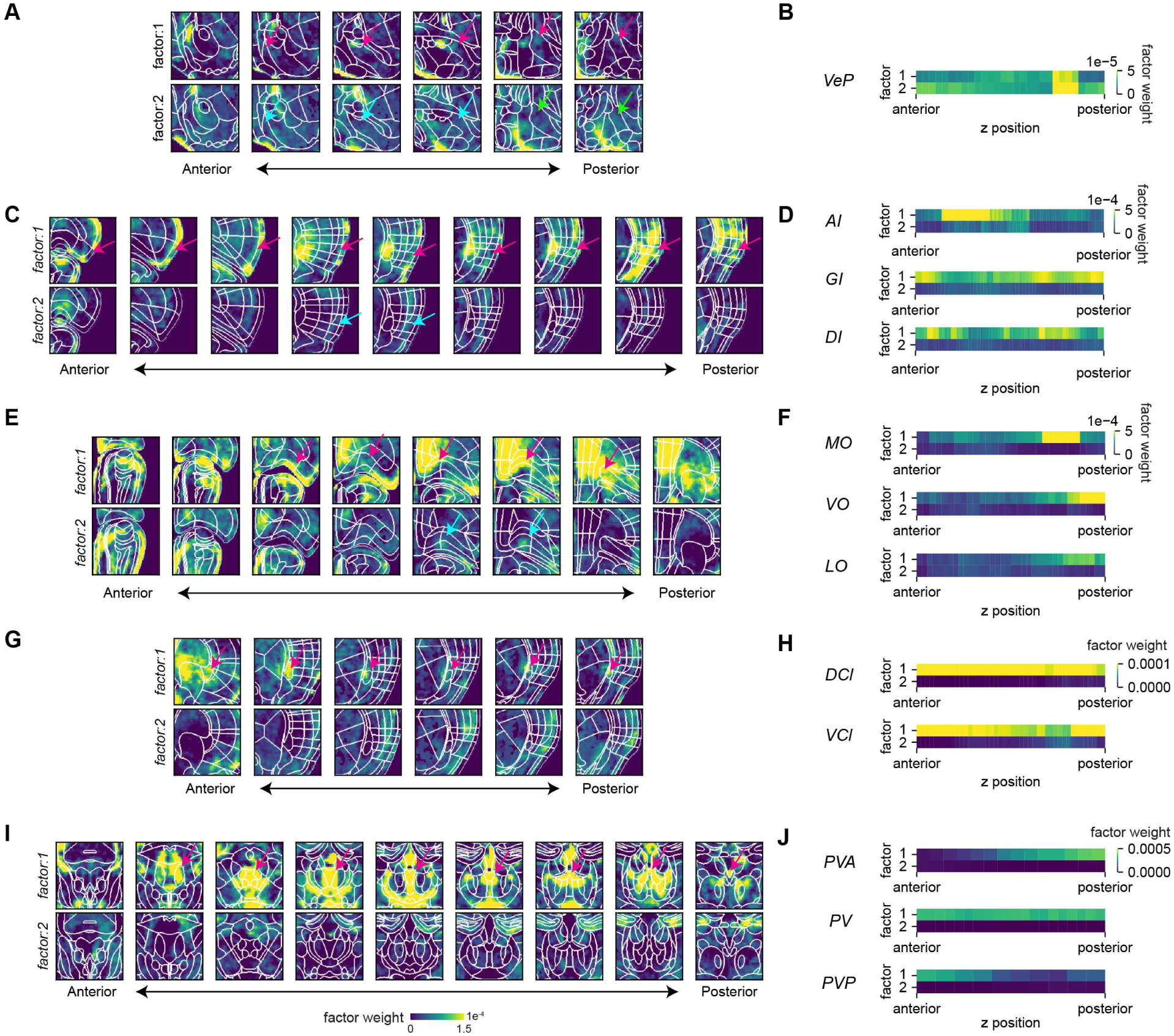
Distribution of morphine responsive factors, related to Figure 5. (A**–**J). Representative 50-μm coronal planes showing factor weights for factor 1 and factor 2 in the ventral pallidum (A), insular cortex (C), orbitofrontal cortex (E), claustrum (G) and paraventricular thalamic nucleus (I) along the anterior-posterior axis. Heatmap showing average factor probability in the ventral pallidum (VeP) (B), insular cortex (D), orbitofrontal cortex (F), claustrum (H) and paraventricular thalamic nucleus (J) along the anterior-posterior axis. Insular cortex GI: granular layer, DI: dysgranular layer, AI: agranular layer. Orbitofrontal cortex MO: medial region, VO: ventral region, LO: lateral region. Claustrum DCl: dorsal region, VCl: ventral region. Paraventricular thalamic nucleus PVA: anterior part, PV: intermediate part, PVP: posterior part.

**Figure S8.**
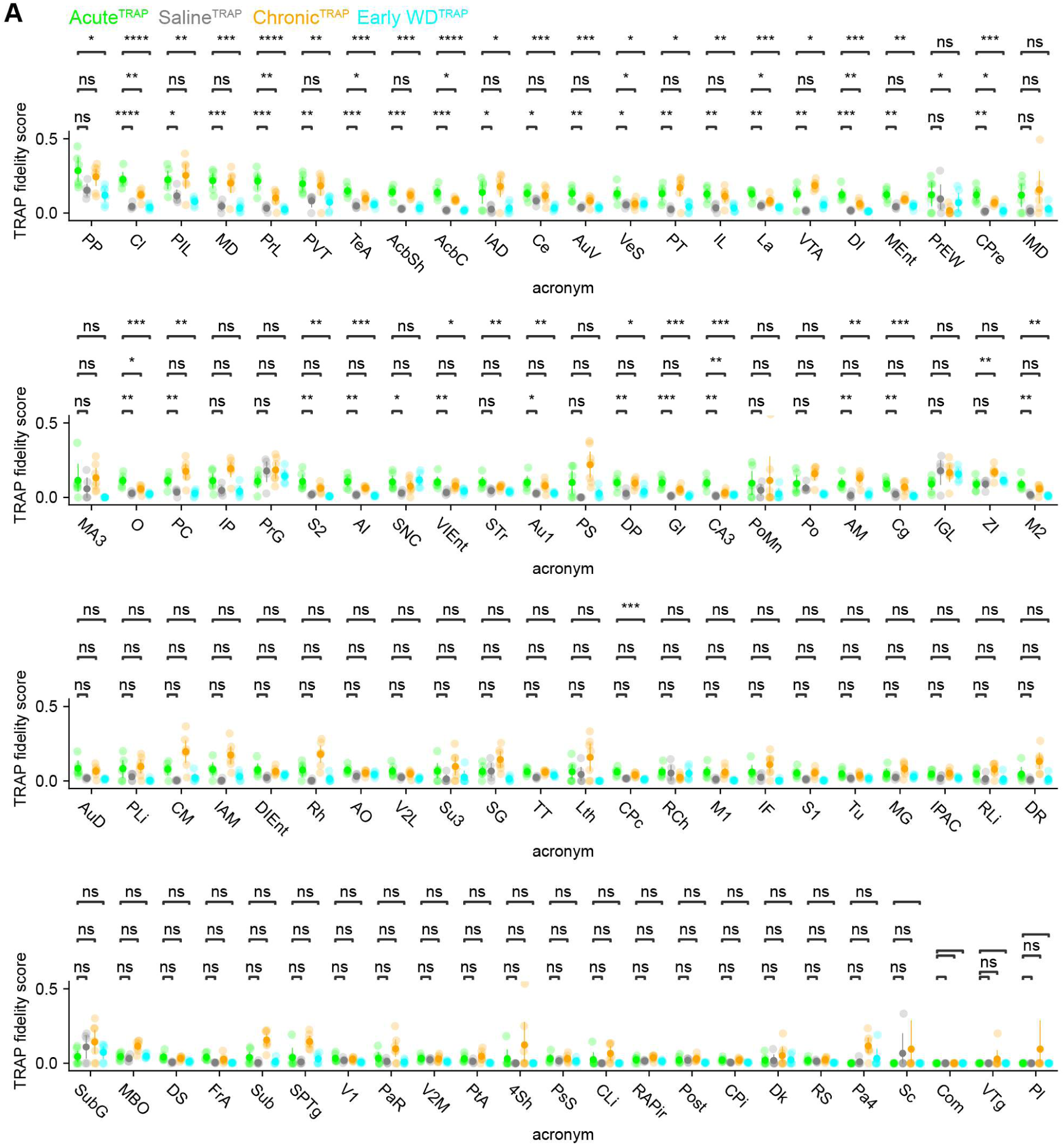
Analysis of acute morphine specific neural ensemble, related to Figure 6. Scatter plot showing TRAP fidelity score in 89 brain regions identified by BRANCH in Figure 2. Student’s t-test was conducted between Acute^TRAP^ and every other condition with post-hoc Benjamini/Hochberg correction. Additional statistics are shown in **Table 6**.

**Figure S9.**
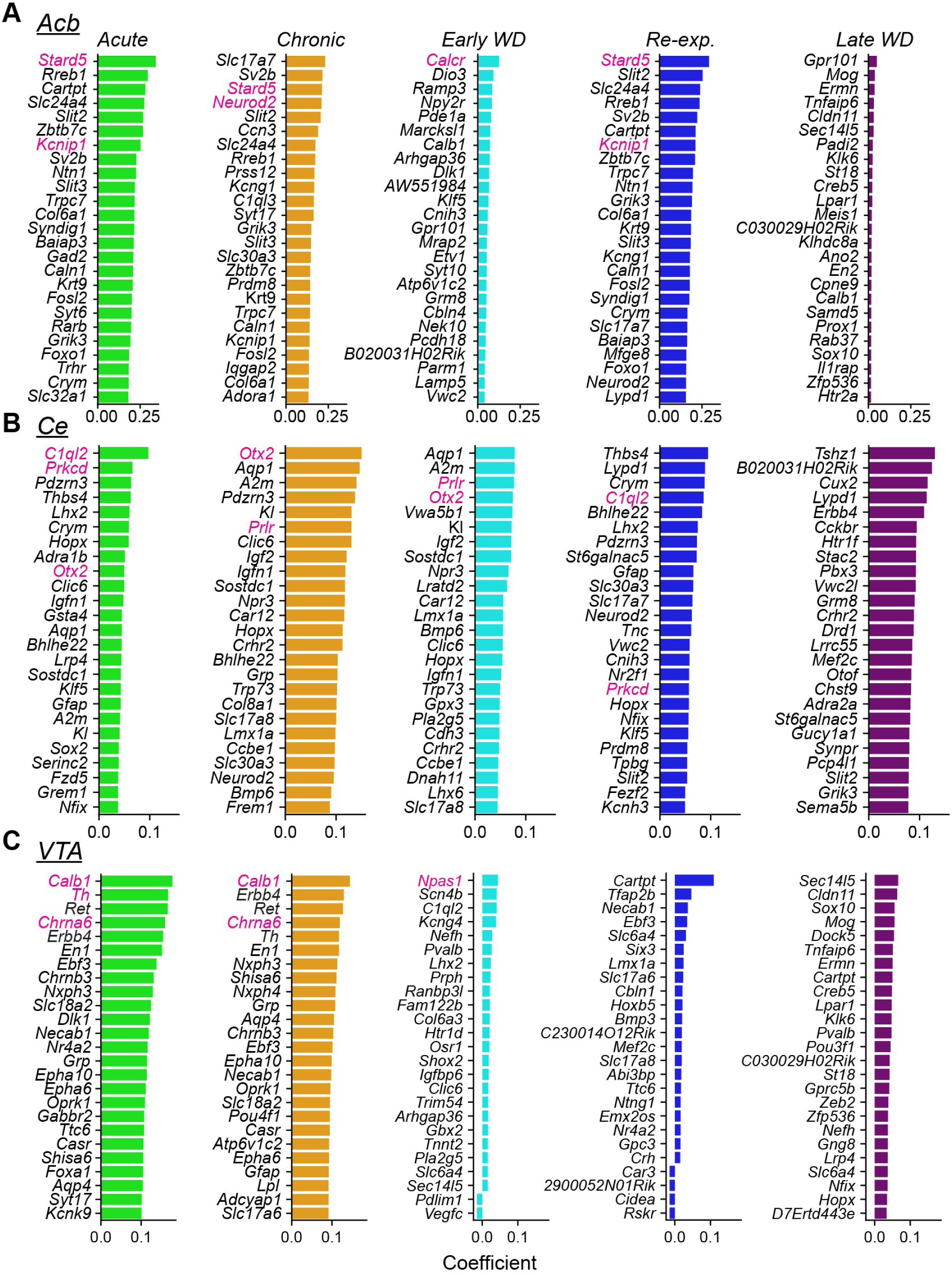
Identification of key molecular markers using c-Fos+ predictive activity map, related to Figure 7. (A**–**C) Bar plot showing the top 30 genes that were significantly correlated with the highest Pearsons coefficient values for each drug condition in the (A) Acb, (B) Ce and (C) VTA (alpha = 0.05 after Benjamini/Hochberg correction). Genes that are shown in Figure 7 are highlighted in magenta. Additional statistics are shown in **Table 7**.

**Figure S10.**
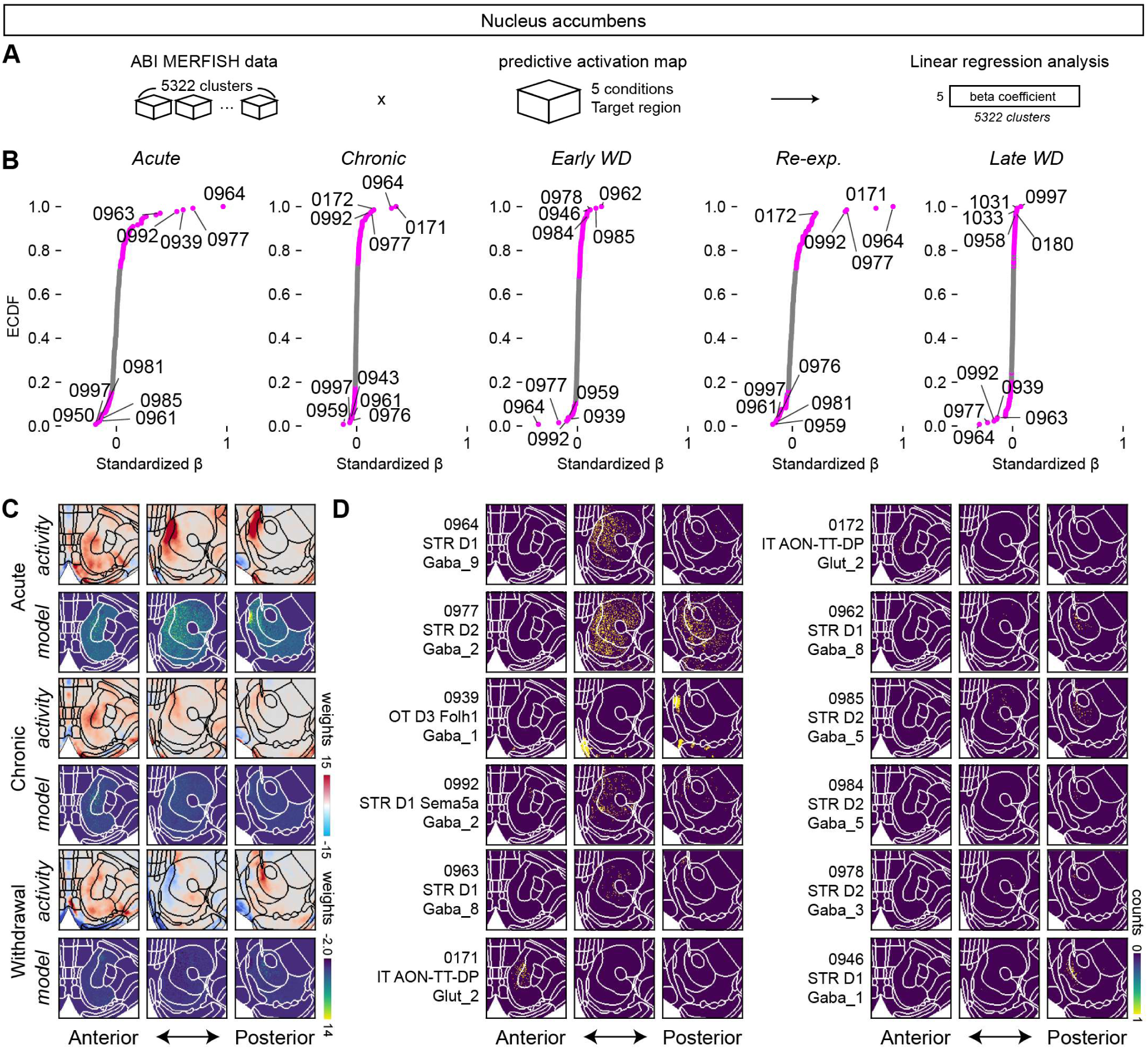
Identification of key cell types using c-Fos+ predictive activity map in the nucleus accumbens, related to Figure 7. (A) Schematic overview of the regression framework. Cell-type cell count maps derived from the ABI MERFISH dataset (5,322 clusters) were used as predictors to explain voxelwise predictive activity maps across five morphine-related conditions in the nucleus accumbens. Linear regression coefficients were estimated for each cluster and condition. (B) Empirical cumulative distribution functions (ECDFs) of standardized regression coefficients (β) for acute, chronic, earlyWD, re-exposure, and lateWD conditions. Individual clusters are labeled by their four-digit IDs; clusters with permutation p-values after FDR correction (Benjamini–Hochberg, alpha = 0.05) are shown in magenta. Additional statistics are shown in **Table 8**. (C) Representative coronal planes showing observed predictive activity maps (top row, “activity”) and regression-modeled predictions (bottom row, “model”) for acute, chronic, and withdrawal conditions. (D) Spatial distributions of representative clusters with high-magnitude coefficients across conditions.

**Figure S11.**
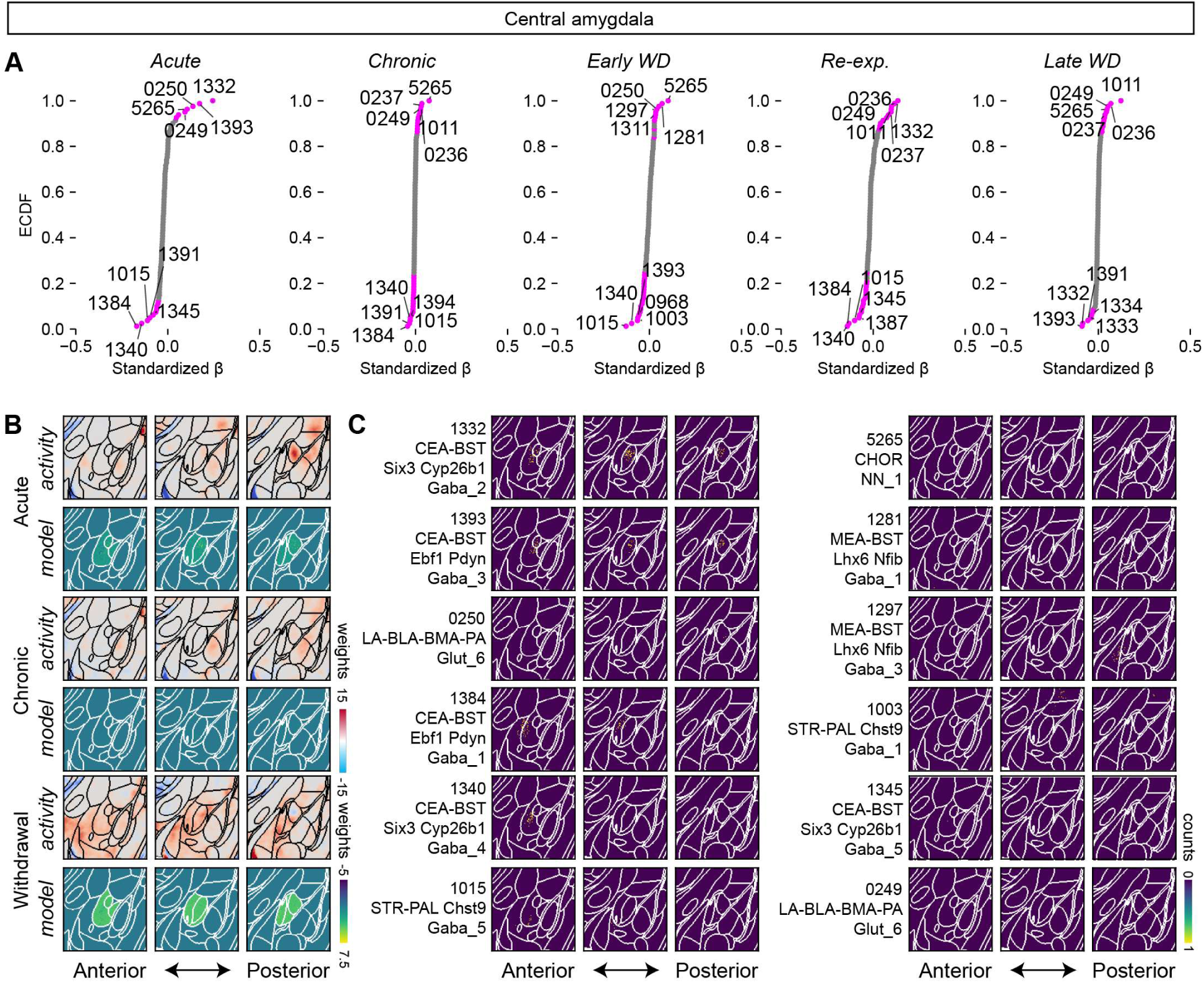
Identification of key cell types using c-Fos+ predictive activity map in the central amygdala, related to Figure 7. (A) Empirical cumulative distribution functions (ECDFs) of standardized regression coefficients (β) for acute, chronic, earlyWD, re-exposure, and lateWD conditions. Individual clusters are labeled by their four-digit IDs; clusters with permutation p-values after FDR correction (Benjamini–Hochberg, α = 0.05) are shown in magenta. Additional statistics are shown in **Table 8**. (B) Representative coronal planes showing observed predictive activity maps (top row, “activity”) and regression-modeled predictions (bottom row, “model”) for acute, chronic, and withdrawal conditions. (C) Spatial distributions of representative clusters with high-magnitude coefficients across conditions.

**Figure S12.**
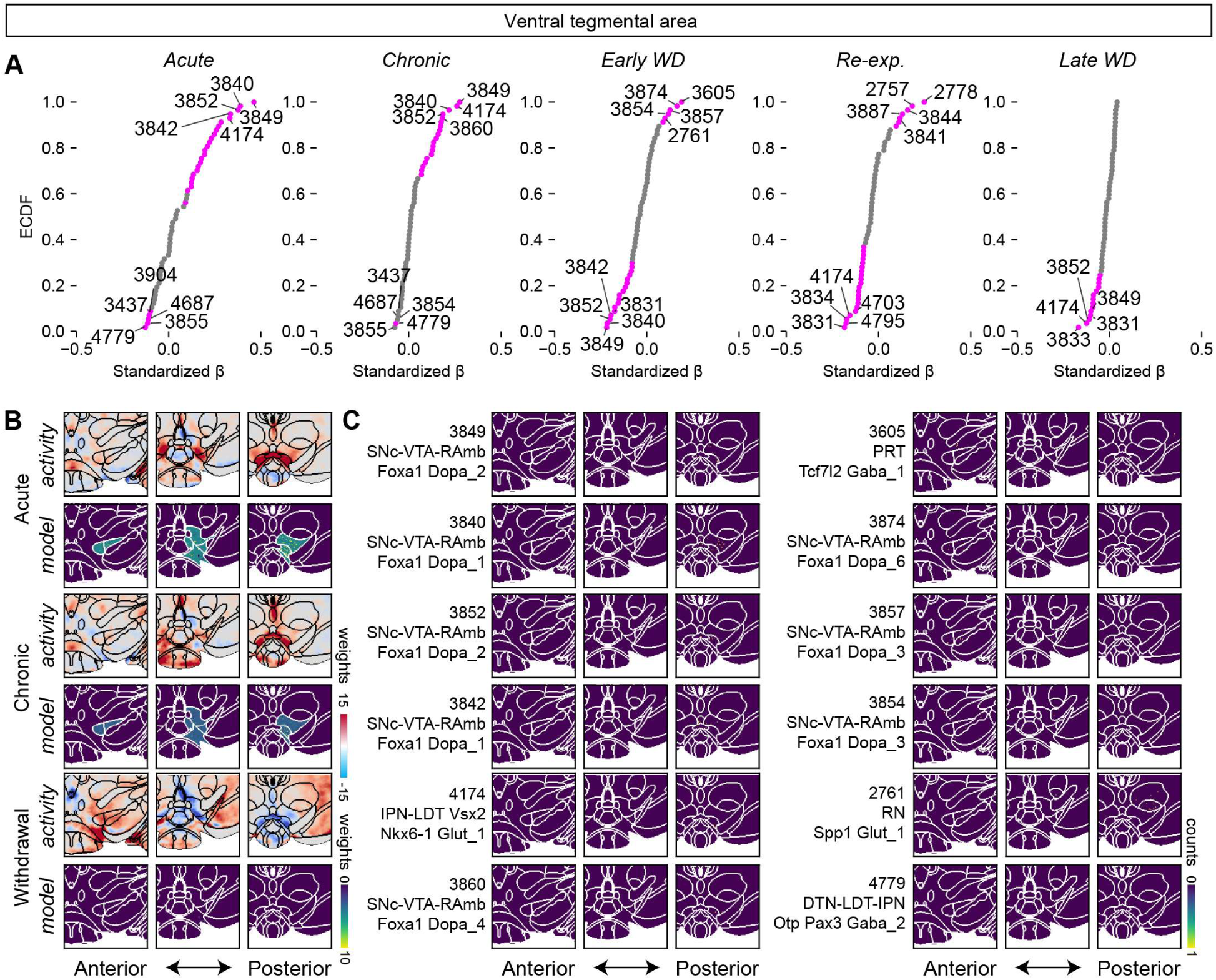
Identification of key cell types using c-Fos+ predictive activity map in the ventral tegmental area, related to Figure 7. (A) Empirical cumulative distribution functions (ECDFs) of standardized regression coefficients (β) for acute, chronic, earlyWD, re-exposure, and lateWD conditions. Individual clusters are labeled by their four-digit IDs; clusters with permutation p-values after FDR correction (Benjamini–Hochberg, α = 0.05) are shown in magenta. Additional statistics are shown in **Table 8**. (B) Representative coronal planes showing observed predictive activity maps (top row, “activity”) and regression-modeled predictions (bottom row, “model”) for acute, chronic, and withdrawal conditions. (C) Spatial distributions of representative clusters with high-magnitude coefficients across conditions.

## References

1. Bruchas, M.R., Land, B.B., and Chavkin, C. (2010). The dynorphin/kappa opioid system as a modulator of stress-induced and pro-addictive behaviors. Brain Research 1314, 44–55. 10.1016/j.brainres.2009.08.062.

2. Basbaum, A.I., Bautista, D.M., Scherrer, G., and Julius, D. (2009). Cellular and Molecular Mechanisms of Pain. Cell 139, 267–284. 10.1016/j.cell.2009.09.028.

3. Fields, H.L., and Margolis, E.B. (2015). Understanding opioid reward. Trends in Neurosciences 38, 217–225. 10.1016/j.tins.2015.01.002.

4. Lüscher, C. (2016). The Emergence of a Circuit Model for Addiction. Annu. Rev. Neurosci. 39, 257–276. 10.1146/annurev-neuro-070815-013920.

5. George, O., and Koob, G.F. (2017). Individual differences in the neuropsychopathology of addiction. Dialogues in Clinical Neuroscience 19, 217–229. 10.31887/DCNS.2017.19.3/gkoob.

6. Koob, G. (2001). Drug Addiction, Dysregulation of Reward, and Allostasis. Neuropsychopharmacology 24, 97–129. 10.1016/S0893-133X(00)00195-0.

7. Reeves, K.C., Shah, N., Muñoz, B., and Atwood, B.K. (2022). Opioid Receptor-Mediated Regulation of Neurotransmission in the Brain. Front. Mol. Neurosci. 15, 919773. 10.3389/fnmol.2022.919773.

8. Peciña, S., Smith, K.S., and Berridge, K.C. (2006). Hedonic Hot Spots in the Brain. Neuroscientist 12, 500–511. 10.1177/1073858406293154.

9. Morales, I., and Berridge, K.C. (2020). ‘Liking’ and ‘wanting’ in eating and food reward: Brain mechanisms and clinical implications. Physiology & Behavior 227, 113152. 10.1016/j.physbeh.2020.113152.

10. Peciña, S. (2008). Opioid reward ‘liking’ and ‘wanting’ in the nucleus accumbens. Physiology & Behavior 94, 675–680. 10.1016/j.physbeh.2008.04.006.

11. Berridge, K.C., and Kringelbach, M.L. (2015). Pleasure Systems in the Brain. Neuron 86, 646–664. 10.1016/j.neuron.2015.02.018.

12. Peciña, S., and Berridge, K.C. (2005). Hedonic Hot Spot in Nucleus Accumbens Shell: Where Do μ-Opioids Cause Increased Hedonic Impact of Sweetness? J. Neurosci. 25, 11777–11786. 10.1523/JNEUROSCI.2329-05.2005.

13. Smith, K.S., and Berridge, K.C. (2005). The Ventral Pallidum and Hedonic Reward: Neurochemical Maps of Sucrose “Liking” and Food Intake. J. Neurosci. 25, 8637–8649. 10.1523/JNEUROSCI.1902-05.2005.

14. Castro, D.C., and Berridge, K.C. (2014). Opioid Hedonic Hotspot in Nucleus Accumbens Shell: Mu, Delta, and Kappa Maps for Enhancement of Sweetness “Liking” and “Wanting.” J. Neurosci. 34, 4239–4250. 10.1523/JNEUROSCI.4458-13.2014.

15. Franceschini, A., Costantini, I., Pavone, F.S., and Silvestri, L. (2020). Dissecting Neuronal Activation on a Brain-Wide Scale With Immediate Early Genes. Front. Neurosci. 14, 569517. 10.3389/fnins.2020.569517.

16. Ueda, H.R., Ertürk, A., Chung, K., Gradinaru, V., Chédotal, A., Tomancak, P., and Keller, P.J. (2020). Tissue clearing and its applications in neuroscience. Nat Rev Neurosci 21, 61–79. 10.1038/s41583-019-0250-1.

17. Hubbard, E., Derdeyn, P., Galinato, V.M., Wu, A., Bartas, K., Mahler, S.V., and Beier, K.T. (2025). Neural basis of adolescent THC-induced potentiation of opioid responses later in life. Neuropsychopharmacol. 50, 818–827. 10.1038/s41386-024-02033-8.

18. Vasylieva, I., Smith, M.C., Aravind, E., He, K., Ling, T., Kozel, J., Puig, S., Kedziora, K.M., Scarlett, J.J., Joseph, P.N., et al. (2025). Brain-wide mapping reveals temporal and sexually dimorphic opioid actions. Preprint, 10.1101/2025.02.19.638902 https://doi.org/10.1101/2025.02.19.638902.

19. Tan, B., Browne, C.J., Nöbauer, T., Vaziri, A., Friedman, J.M., and Nestler, E.J. (2024). Drugs of abuse hijack a mesolimbic pathway that processes homeostatic need. Science 384, eadk6742. 10.1126/science.adk6742.

20. Chaudun, F., Python, L., Liu, Y., Hiver, A., Cand, J., Kieffer, B.L., Valjent, E., and Lüscher, C. (2024). Distinct µ-opioid ensembles trigger positive and negative fentanyl reinforcement. Nature 630, 141–148. 10.1038/s41586-024-07440-x.

21. Keyes, P.C., Adams, E.L., Chen, Z., Bi, L., Nachtrab, G., Wang, V.J., Tessier-Lavigne, M., Zhu, Y., and Chen, X. (2020). Orchestrating Opiate-Associated Memories in Thalamic Circuits. Neuron 107, 1113–1123.e4. 10.1016/j.neuron.2020.06.028.

22. Smith, A.C.W., Ghoshal, S., Centanni, S.W., Heyer, M.P., Corona, A., Wills, L., Andraka, E., Lei, Y., O’Connor, R.M., Caligiuri, S.P.B., et al. (2024). A master regulator of opioid reward in the ventral prefrontal cortex. Science 384. 10.1126/science.adn0886.

23. Park, Y.-G., Sohn, C.H., Chen, R., McCue, M., Yun, D.H., Drummond, G.T., Ku, T., Evans, N.B., Oak, H.C., Trieu, W., et al. (2019). Protection of tissue physicochemical properties using polyfunctional crosslinkers. Nature Biotechnology 37, 73–83. 10.1038/nbt.4281.

24. Chon, U., Vanselow, D.J., Cheng, K.C., and Kim, Y. (2019). Enhanced and unified anatomical labeling for a common mouse brain atlas. Nature Communications 10. 10.1038/s41467-019-13057-w.

25. Bogomolov, M., Peterson, C.B., Benjamini, Y., and Sabatti, C. (2021). Hypotheses on a tree: new error rates and testing strategies. Biometrika 108, 575–590. 10.1093/biomet/asaa086.

26. Bontempi, B., and Sharp, F.R. Systemic Morphine-Induced Fos Protein in the Rat Striatum and Nucleus Accumbens Is Regulated by ␮ Opioid Receptors in the Substantia Nigra and Ventral Tegmental Area.

27. Liu, J., Nickolenko, J., and Sharp, F.R. (1994). Morphine induces c-fos and junB in striatum and nucleus accumbens via D1 and N-methyl-D-aspartate receptors. Proc. Natl. Acad. Sci. U.S.A. 91, 8537–8541. 10.1073/pnas.91.18.8537.

28. Lammel, S., Lim, B.K., Ran, C., Huang, K.W., Betley, M.J., Tye, K.M., Deisseroth, K., and Malenka, R.C. (2012). Input-specific control of reward and aversion in the ventral tegmental area. Nature 491, 212–217. 10.1038/nature11527.

29. Matsui, A., Jarvie, B.C., Robinson, B.G., Hentges, S.T., and Williams, J.T. (2014). Separate GABA Afferents to Dopamine Neurons Mediate Acute Action of Opioids, Development of Tolerance, and Expression of Withdrawal. Neuron 82, 1346–1356. 10.1016/j.neuron.2014.04.030.

30. Tan, K.R., Yvon, C., Turiault, M., Mirzabekov, J.J., Doehner, J., Labouèbe, G., Deisseroth, K., Tye, K.M., and Lüscher, C. (2012). GABA Neurons of the VTA Drive Conditioned Place Aversion. Neuron 73, 1173–1183. 10.1016/j.neuron.2012.02.015.

31. Johnson, S.W., and North, R.A. (1992). Two types of neurone in the rat ventral tegmental area and their synaptic inputs. The Journal of Physiology 450, 455–468. 10.1113/jphysiol.1992.sp019136.

32. Koob, G.F. (2020). Neurobiology of Opioid Addiction: Opponent Process, Hyperkatifeia, and Negative Reinforcement. Biological Psychiatry 87, 44–53. 10.1016/j.biopsych.2019.05.023.

33. Fox, A.S., Oler, J.A., Tromp, D.P.M., Fudge, J.L., and Kalin, N.H. (2015). Extending the amygdala in theories of threat processing. Trends in Neurosciences 38, 319–329. 10.1016/j.tins.2015.03.002.

34. Neugebauer, V., Li, W., Bird, G.C., and Han, J.S. (2004). The Amygdala and Persistent Pain. Neuroscientist 10, 221–234. 10.1177/1073858403261077.

35. Zhang, W.-H., Zhang, J.-Y., Holmes, A., and Pan, B.-X. (2021). Amygdala Circuit Substrates for Stress Adaptation and Adversity. Biological Psychiatry 89, 847–856. 10.1016/j.biopsych.2020.12.026.

36. Kaplan, G.B., and Thompson, B.L. (2023). Neuroplasticity of the extended amygdala in opioid withdrawal and prolonged opioid abstinence. Front. Pharmacol. 14, 1253736. 10.3389/fphar.2023.1253736.

37. McInnes, L., Healy, J., and Melville, J. (2020). UMAP: Uniform Manifold Approximation and Projection for Dimension Reduction. Preprint at arXiv, 10.48550/arXiv.1802.03426 https://doi.org/10.48550/arXiv.1802.03426.

38. Friedmann, D., Gonzalez, X., Moses, A., Watts, T., Degleris, A., Ticea, N., Song, J.H., Datta, S.R., Linderman, S.W., and Luo, L. (2024). Concerted modulation of spontaneous behavior and time-integrated whole-brain neuronal activity by serotonin receptors. Preprint, 10.1101/2024.08.02.606282 https://doi.org/10.1101/2024.08.02.606282.

39. Castro, D.C., and Berridge, K.C. (2017). Opioid and orexin hedonic hotspots in rat orbitofrontal cortex and insula. Proc. Natl. Acad. Sci. U.S.A. 114. 10.1073/pnas.1705753114.

40. Allen, W.E., DeNardo, L.A., Chen, M.Z., Liu, C.D., Loh, K.M., Fenno, L.E., Ramakrishnan, C., Deisseroth, K., and Luo, L. (2017). Thirst-associated preoptic neurons encode an aversive motivational drive. Science 357, 1149–1155. 10.1126/science.aan6747.

41. Zhang, M., Pan, X., Jung, W., Halpern, A.R., Eichhorn, S.W., Lei, Z., Cohen, L., Smith, K.A., Tasic, B., Yao, Z., et al. (2023). Molecularly defined and spatially resolved cell atlas of the whole mouse brain. Nature 624, 343–354. 10.1038/s41586-023-06808-9.

42. Haubensak, W., Kunwar, P.S., Cai, H., Ciocchi, S., Wall, N.R., Ponnusamy, R., Biag, J., Dong, H.-W., Deisseroth, K., Callaway, E.M., et al. (2010). Genetic dissection of an amygdala microcircuit that gates conditioned fear. Nature 468, 270–276. 10.1038/nature09553.

43. Yao, Z., Van Velthoven, C.T.J., Kunst, M., Zhang, M., McMillen, D., Lee, C., Jung, W., Goldy, J., Abdelhak, A., Aitken, M., et al. (2023). A high-resolution transcriptomic and spatial atlas of cell types in the whole mouse brain. Nature 624, 317–332. 10.1038/s41586-023-06812-z.

44. Choi, H.M.T., Beck, V.A., and Pierce, N.A. (2014). Next-Generation in Situ Hybridization Chain Reaction: Higher Gain, Lower Cost, Greater Durability. ACS Nano 8, 4284–4294. 10.1021/nn405717p.

45. Castro, D.C., Oswell, C.S., Zhang, E.T., Pedersen, C.E., Piantadosi, S.C., Rossi, M.A., Hunker, A.C., Guglin, A., Morón, J.A., Zweifel, L.S., et al. (2021). An endogenous opioid circuit determines state-dependent reward consumption. Nature 598, 646–651. 10.1038/s41586-021-04013-0.

46. Welsch, L., Colantonio, E., Frison, M., Johnson, D.A., McClain, S.P., Mathis, V., Banghart, M.R., Ben Hamida, S., Darcq, E., and Kieffer, B.L. (2023). Mu Opioid Receptor–Expressing Neurons in the Dorsal Raphe Nucleus Are Involved in Reward Processing and Affective Behaviors. Biological Psychiatry 94, 842–851. 10.1016/j.biopsych.2023.05.019.

47. Qing-Ping, W., and Nakai, Y. (1994). The dorsal raphe: An important nucleus in pain modulation. Brain Research Bulletin 34, 575–585. 10.1016/0361-9230(94)90143-0.

48. Tao, R., and Auerbach, S.B. (2005). μ-Opioids disinhibit and κ-opioids inhibit serotonin efflux in the dorsal raphe nucleus. Brain Research 1049, 70–79. 10.1016/j.brainres.2005.04.076.

49. Alvarez-Bagnarol, Y., García, R., Vendruscolo, L.F., and Morales, M. (2023). Inhibition of dorsal raphe GABAergic neurons blocks hyperalgesia during heroin withdrawal. Neuropsychopharmacol. 48, 1300–1308. 10.1038/s41386-023-01620-5.

50. Pan, Z.Z., Williams, J.T., and Osborne, P.B. (1990). Opioid actions on single nucleus raphe magnus neurons from rat and guinea-pig in vitro. The Journal of Physiology 427, 519–532. 10.1113/jphysiol.1990.sp018185.

51. Coutens, B., and Ingram, S.L. (2023). Key differences in regulation of opioid receptors localized to presynaptic terminals compared to somas: Relevance for novel therapeutics. Neuropharmacology 226, 109408. 10.1016/j.neuropharm.2022.109408.

52. Lutz, P.-E., and Kieffer, B.L. (2013). Opioid receptors: distinct roles in mood disorders. Trends in Neurosciences 36, 195–206. 10.1016/j.tins.2012.11.002.

53. Bals-Kubik, R., Herz, A., and Shippenberg, T.S. (1989). Evidence that the aversive effects of opioid antagonists and ?-agonists are centrally mediated. Psychopharmacology 98, 203–206. 10.1007/bf00444692.

54. Land, B.B., Bruchas, M.R., Lemos, J.C., Xu, M., Melief, E.J., and Chavkin, C. (2008). The Dysphoric Component of Stress Is Encoded by Activation of the Dynorphin κ-Opioid System. J. Neurosci. 28, 407–414. 10.1523/jneurosci.4458-07.2008.

55. Cahill, C.M., Taylor, A.M.W., Cook, C., Ong, E., MorÃ^3^n, J.A., and Evans, C.J. (2014). Does the kappa opioid receptor system contribute to pain aversion? Front. Pharmacol. 5. 10.3389/fphar.2014.00253.

56. Yang, D., Wang, Y., Qi, T., Zhang, X., Shen, L., Ma, J., Pang, Z., Lal, N.K., McClatchy, D.B., Seradj, S.H., et al. (2024). Phosphorylation of pyruvate dehydrogenase inversely associates with neuronal activity. Neuron 112, 959–971.e8. 10.1016/j.neuron.2023.12.015.

57. DeNardo, L.A., Liu, C.D., Allen, W.E., Adams, E.L., Friedmann, D., Fu, L., Guenthner, C.J., Tessier-Lavigne, M., and Luo, L. (2019). Temporal evolution of cortical ensembles promoting remote memory retrieval. Nat Neurosci 22, 460–469. 10.1038/s41593-018-0318-7.

58. Cai, X., Huang, H., Kuzirian, M.S., Snyder, L.M., Matsushita, M., Lee, M.C., Ferguson, C., Homanics, G.E., Barth, A.L., and Ross, S.E. (2016). Generation of a KOR-Cre knockin mouse strain to study cells involved in kappa opioid signaling. Genesis 54, 29–37. 10.1002/dvg.22910.

59. Ishii, K.K., Hashikawa, K., Chea, J., Yin, S., Fox, R.E., Kan, S., Shah, M., Zhou, Z.C., Navarrete, J., Murry, A.D., et al. (2025). Post-ejaculatory inhibition of female sexual drive via heterogeneous neuronal ensembles in the medial preoptic area. eLife 12, RP91765. 10.7554/eLife.91765.

60. Stringer, C., Wang, T., Michaelos, M., and Pachitariu, M. (2021). Cellpose: a generalist algorithm for cellular segmentation. Nature Methods 18, 100–106. 10.1038/s41592-020-01018-x.

61. Imbert, A., Ouyang, W., Safieddine, A., Coleno, E., Zimmer, C., Bertrand, E., Walter, T., and Mueller, F. (2022). FISH-quant v2: a scalable and modular tool for smFISH image analysis. RNA 28, 786–795. 10.1261/rna.079073.121.

62. Berg, S., Kutra, D., Kroeger, T., Straehle, C.N., Kausler, B.X., Haubold, C., Schiegg, M., Ales, J., Beier, T., Rudy, M., et al. (2019). ilastik: interactive machine learning for (bio)image analysis. Nature Methods 16, 1226–1232. 10.1038/s41592-019-0582-9.

63. Renier, N., Adams, E.L., Kirst, C., Wu, Z., Azevedo, R., Kohl, J., Autry, A.E., Kadiri, L., Umadevi Venkataraju, K., Zhou, Y., et al. (2016). Mapping of Brain Activity by Automated Volume Analysis of Immediate Early Genes. Cell 165, 1789–1802. 10.1016/j.cell.2016.05.007.

64. Boerner, T.J., Deems, S., Furlani, T.R., Knuth, S.L., and Towns, J. (2023). ACCESS: Advancing Innovation: NSF’s Advanced Cyberinfrastructure Coordination Ecosystem: Services & Support. In Practice and Experience in Advanced Research Computing (ACM), pp. 173–176. 10.1145/3569951.3597559.

65. Brown, S.T., Buitrago, P., Hanna, E., Sanielevici, S., Scibek, R., and Nystrom, N.A. (2021). Bridges-2: A Platform for Rapidly-Evolving and Data Intensive Research. In Practice and Experience in Advanced Research Computing (ACM), pp. 1–4. 10.1145/3437359.3465593.

66. Wang, Q., Ding, S.-L., Li, Y., Royall, J., Feng, D., Lesnar, P., Graddis, N., Naeemi, M., Facer, B., Ho, A., et al. (2020). The Allen Mouse Brain Common Coordinate Framework: A 3D Reference Atlas. Cell 181, 936–953.e20. 10.1016/j.cell.2020.04.007.

67. Simes, R.J. (1986). An improved Bonferroni procedure for multiple tests of significance. Biometrika 73, 751–754. 10.1093/biomet/73.3.751.

68. Benjamini, Y., and Hochberg, Y. (1995). Controlling the False Discovery Rate: A Practical and Powerful Approach to Multiple Testing. Journal of the Royal Statistical Society Series B: Statistical Methodology 57, 289–300. 10.1111/j.2517-6161.1995.tb02031.x.

